# Astrocytic Nonsense-mediated mRNA decay regulates calcium signaling to support synapse function and influence anxiety-like behavior

**DOI:** 10.1101/2025.11.18.689044

**Authors:** Pablo J. Lituma, Aykut Deveci, Estibaliz Barrio-Alonso, Kun Tan, Miles F. Wilkinson, Pablo E. Castillo, Dilek Colak

## Abstract

How astrocytes achieve their diverse roles in the brain at the molecular level is poorly understood. In this study, we leverage mouse models, electrophysiology, calcium imaging, behavioral assays, and bioinformatic approaches to demonstrate that astrocyte activity and astrocyte-mediated mouse behavior depends on the highly conserved and selective RNA turnover pathway-nonsense-mediated RNA decay (NMD). Conditional deletion of the core NMD gene, *Upf2*, in mature astrocytes leads to enhanced basal Ca^2+^ signaling coupled with synapse dysfunction and elevated anxiety-like behavior. Restoring basal Ca^2+^ signaling in NMD-deficient astrocytes shifted synaptic transmission and minimized anxiety-associated behavior. Molecular bioinformatic analysis identified specific NMD target transcripts in astrocytes as candidates influencing calcium signaling pathways and neuro-glia interactions that support brain function. Our study is the first to demonstrate specific roles for NMD in astrocytes.

## INTRODUCTION

Astrocytes are highly communicative. A single mouse cortical astrocyte can contact ∼100,000 synapses (1). This number has been suggested to reach up to 2,000,000 synapses for a human astrocyte (2). Thus, astroglia cells are in powerful position to monitor and respond to neural circuit dynamics. Astrocytes detect synaptic activity via membrane transporters and receptors that elevate intracellular Ca^2+^ levels as a means to activate downstream signaling pathways that then trigger release of gliotransmitters and other molecules to modulate neuronal excitability and synaptic function (3–6). Astrocytes also have other functions, including synapse engulfment and autophagy, both of which are critical for neural circuit homeostasis, and, if faulty, contribute to neurodegenerative brain states in adulthood (7, 8).

The molecular pathways underlying astrocyte’s diverse functions are poorly understood. An attractive molecular mechanism that has the potential to have a role in spatially- and temporally-restricted activities in astrocytes is post-transcriptional regulation. Deficits in post-transcriptional regulation in neurons are known to result in ectopic protein synthesis and lead to developmental and cognitive disabilities (9–15). Meanwhile, the impact of disrupted post-transcriptional regulation influencing mRNA stability in astrocytes and other brain cell-types remains to be elucidated (16, 17). The best characterized RNA turnover pathway is nonsense-mediated RNA decay (NMD) (18–20). This highly conserved and selective RNA degradation mechanism was first discovered because of its ability to degrade aberrant mRNAs harboring mutations that create premature termination codons (PTCs). NMD was subsequently found to increase the decay rate of subsets of normal mRNAs (21). These normal mRNAs harbor “NMD-inducing features”—such as an exon-exon junction downstream of the main open reading frame (ORF) and long 3’ untranslated regions (UTRs)—that trigger their decay via NMD (21–23). The discovery that NMD destabilizes many normal mRNAs has led to the hypothesis that NMD regulates normal biological events, a hypothesis that has been supported by a plethora of studies showing that NMD influences numerous fundamental processes, including development, differentiation, cell cycle regulation, cell survival, stress response, and autophagy (21).

NMD is a complex pathway that depends on many protein factors, including the core NMD factors UPF1, UPF2, and UPF3B (24). In humans, mutations in *UPF3B* have been shown to cause intellectual disability (25–27) and individuals with *UPF3B* mutations often suffer from autism, schizophrenia, and attention-deficit/hyperactivity disorder (25–29), highlighting the crucial role of regulated RNA turnover in the nervous system (25). Similarly, mutations or copy number variants (CNVs) of NMD machinery genes *UPF2*, *UPF3a*, *SMG6* or null mutations of *UPF3B* are associated with neurodevelopmental and neuropsychiatric disorders (21, 24, 25, 27–30). Given that NMD is ubiquitous and NMD target mRNAs are tissue-, cell type-, and NMD factor-specific (31) how NMD dysfunction precisely contributes to brain disorders in humans is unknown. Nonetheless, NMD perturbation via *Upf2* manipulation in neurons using transgenic mice revealed impairments to cell-cycle regulation, cytoarchitecture, axon guidance, synapse activity, and memory function (22, 23, 32–36). Hence, *Upf2* ablation has proven as a versatile tool to further our understanding of NMD function in mammalian cells (37–41). By leveraging *Upf2* transgenic mouse tools, a comprehensive model of how mRNA degradation impacts synaptic and cognitive function can be constructed by dissecting out the role of NMD in all cell types that modulate these brain processes.

To date, no study has reported on roles for NMD in glial cells. To address this important knowledge gap, we conditionally knocked out the NMD gene, *Upf2*, in astrocytes *in vivo*. Implementing cellular and molecular tools, we reveal that astrocytic UPF2-dependent NMD disruption leads to impaired astrocyte cell size and increased Ca^2+^ activity in hippocampus, visual cortex, and prefrontal cortex. These phenotypes are accompanied with impaired synaptic function and anxiety-like behavior in adult mice. To define a molecular mechanism, using RNA sequencing and an established algorithm that identifies NMD-inducing features in transcripts, we determined the physiological targets of UPF2-dependent NMD in astrocytes. This allowed us to predict the biological pathways regulated by NMD in these cells. NMD targets within astrocytes pointed to morphological remodeling, phagosome formation, and Ca^2+^ signaling. Modulation of Ca^2+^ activity in NMD-deficient astrocytes *in vivo* partially restored cell volume, shifted synaptic transmission and minimized anxiety-like behavior. By characterizing the role of UPF2 dependent-NMD in astrocytes, our study highlights that RNA degradation is a major mechanism that regulates critical astrocytic biological processes required for neuro-glia interactions that ultimately support proper brain function.

## RESULTS

### Conditional deletion of *Upf2* in astrocytes *in vivo* reduces their size

To determine the physiological requirement for NMD in astrocytes, we leveraged mouse transgenic Cre-lox strategies to ablate the *Upf2* gene in adult mice. *Upf2^wt/wt^* and *Upf2^fl/fl^* animals were crossed with the *Aldh1L1-CreER^T2^* transgenic mouse line to provide temporal control of *Upf2* deletion in astrocytes using tamoxifen. The *Aldh1L1-CreER^T2^* transgenic mouse line induces CRE expression in astrocytes with high specificity (42) and is widely implemented to study astrocyte functions. To visualize astrocytes and CRE recombination we co-injected the adeno-associated viruses AAV5-GfaABC1D-Lck-GFP and AAV8-GFAP-mCherry-Cre, respectively, into the hippocampus, visual cortex, and prefrontal cortex of CTRL (*Upf2*^wt/wt^;*Aldh1L1-CreER^T2^*) and cKO (*Upf2*^fl/fl^;*Aldh1L1-CreER^T2^*) mice followed by tamoxifen administration (**Figure 1A**). Immunohistochemistry confirmed the expression of mCherry-Cre and Lck-GFP across the targeted brain areas with negligible neuronal leak (< 1%, **Figure S1**). By anchoring the fluorophore to the membrane, Lck-GFP enables the visualization of fine, ramified branches typically lost in cytosolic imaging, achieving the precision required for morphology analysis. Using spinning disk confocal microscopy, images of Lck-GFP^+^ astrocytes in hippocampus, visual cortex, and prefrontal cortex were acquired (**Figure 1A**). Imaris 3D reconstructions enabled cell volume measurements of CTRL and cKO astrocytes across brain regions (**Figure 1B**). Quantifications revealed that, in all brain regions analyzed, NMD deficiency led to significant reductions in astrocyte cell volume as compared to CTRL (**Figure 1C-D**). Our results suggest that NMD regulates astrocyte morphology and raised the possibility that altered astrocyte cell size may impact neuro-glia interactions in *Upf2* cKO brains.

**Figure 1.**
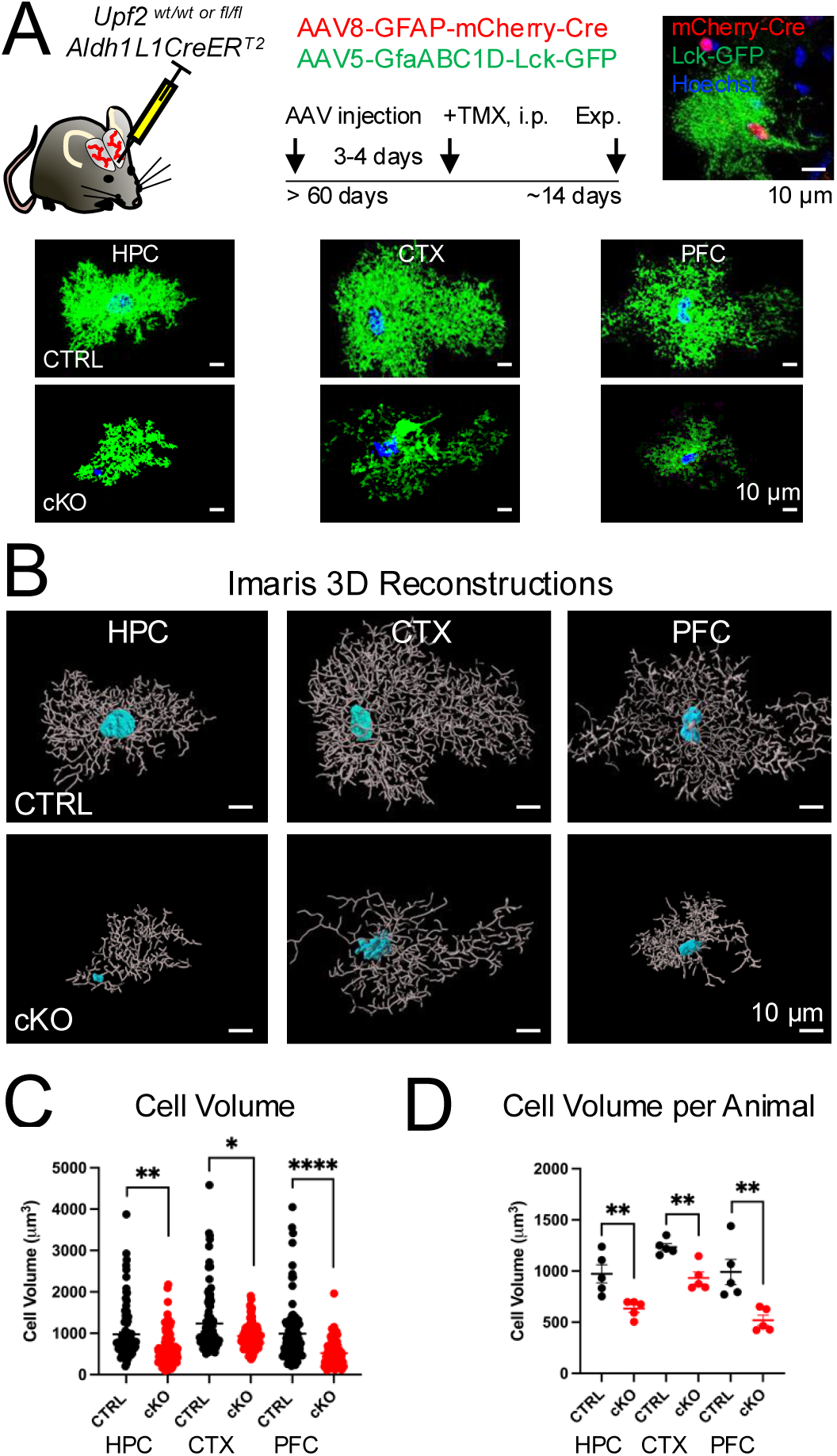
*Upf2* cKO astrocytes show cell size adaptations across brain areas. (**A**) Schematic of adeno-associated viral delivery of AAV8-GFAP-mCherry-Cre and AAV5-GfaABC1D.PI.Lck-GFP.SV40 reporters to hippocampus (HPC), visual cortex (CTX), and prefrontal cortex (PFC). Image of mCherry and Lck-GFP expression in region of interest-ROI. Representative astrocyte morphology images from CTRL *(top)* and *Upf2* cKO *(bottom)* across targeted brain areas. (**B**) Imaris 3D reconstructions of CTRL *(top)* and cKO *(bottom)* astrocytes from HPC, CTX, and PFC. (**C**) *Upf2* mutant astrocytes display reduced cell volume as compared to CTRL in HPC, CTX, and PFC (HPC, CTRL: 973 ± 63, 100 cells; cKO: 634 ± 40, 100 cells, p < 0.001, Mann-Whitney test. CTX, CTRL: 1234 ± 75, 100 cells; cKO: 933 ± 31, 100 cells, p = 0.0216, Mann-Whitney test. PFC, CTRL: 990 ± 72, 100 cells; cKO: 519 ± 30, 100 cells, p < 0.0001, Mann-Whitney test. 20 cells per animal, 5 animals per genotype). (**D**) Cell volume summary data across brain regions per animal analysis (HPC, CTRL: 973 ± 89, cKO: 634 ± 38, p = 0.0079, Mann-Whitney test. CTX, CTRL: 1234 ± 33, cKO: 933 ± 258, p = 0.0079, Mann-Whitney test. PFC, CTRL: 990 ± 124, cKO: 519 ± 50, p = 0.0079, Mann-Whitney test. 5 animals per genotype). Data is presented as mean ± S.E.M. * denotes p < 0.05, ** denotes p < 0.01 **** denotes p < 0.0001. **This figure highlights decreased cell volume in NMD-deficient astrocytes in hippocampus, cortex, and prefrontal cortex.**

### Conditional deletion of *Upf2* in astrocytes *in vivo* disrupts excitatory synapse numbers and spine density

Astrocytes contribute to formation, pruning, and maintenance of synapses in the mammalian central nervous system (7, 43–45). Their unique morphology and interactions with synapses play a crucial role in regulating synapse plasticity and maintenance. Thus, we next sought to determine the impact of astrocytic NMD deficiency on synapse numbers in *Upf2* cKO brains. To facilitate labeling of recombined astrocytes, we employed the Ai6 transgenic mouse line, which drives expression of the ZsGreen1 fluorescent reporter upon CRE recombination. Tamoxifen dosing of *Upf2*^wt/wt^; *Aldh1L1-CreER^T2^; Ai6* (**CTRL**) or *Upf2*^fl/fl^; *Aldh1L1CreER^T2^; Ai6* (**cKO**) triple transgenic mice enabled 488 nm fluorescence of recombined astrocytes (**Figure 2A**). Immunohistochemistry revealed that ∼90% of astrocyte markers GFAP and S100B colocalized with ZsGreen1 in hippocampus and cortex (**Figure S2**) indicating high recombination rate with successful depletion of both UPF2 protein and transcript levels in these cells (**Figure S3**). Moreover, ZsGreen1^+^ *Upf2* cKO astrocytes revealed decreased branch length (**Figure S4**) supporting our morphological observations using the Lck-GFP reporter. To determine the impact of NMD deficiency on excitatory synapse number we quantified PSD-95 levels in ZsGreen1^+^ astrocyte regions of interest (**Figure 2A-B**). Spinning disk confocal microscopy followed by Imaris 3D reconstructions facilitated the quantification of PSD-95^+^ excitatory synapses (**Figure 2B**). Our analysis revealed that *Upf2* cKO brain sections displayed significant reductions in PSD-95 levels across several brain regions when compared to CTRL (**Figure 2C**). The bulk of excitatory synaptic transmission in the brain occurs at dendritic spines. To determine if reduced synapse numbers are accompanied with alterations in spine densities in *Upf2* cKO brains, we labeled excitatory neurons and their spines by injecting adeno-associated virus AAV5-CamKIIα-EGFP to the hippocampus of CTRL (*Upf2*^wt/wt^;*Aldh1L1-CreER^T2^*) and cKO (*Upf2*^fl/fl^;*Aldh1L1-CreER^T2^*) mice followed by tamoxifen administration (**Figure 2D**). Quantifications of spine densities revealed a significant decrease in *Upf2* cKO mice as compared to CTRL (**Figure 2D**). Taken together, these findings support the hypothesis that UPF2-dependent NMD in astrocytes is essential for proper excitatory synapse regulation mediated by these cells.

**Figure 2.**
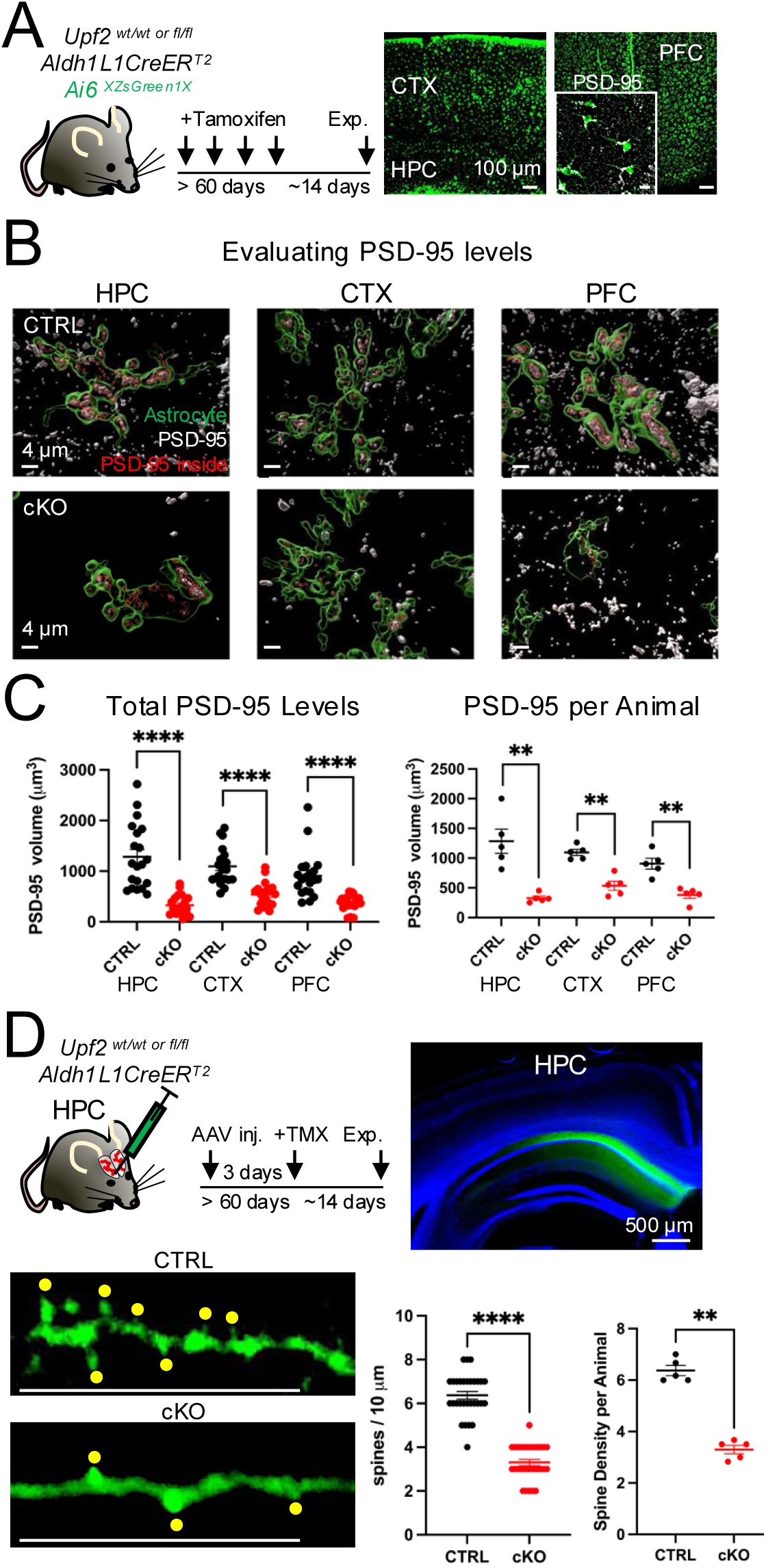
*Upf2* cKO mice display reductions in PSD-95 levels and spine density. (**A**) Schematic of tamoxifen dosing and images of ZsGreen1^+^ astrocytes in CTX, HPC, and PFC. Inset, depicts field of view of ZsGreen1^+^ astrocytes and PSD-95 puncta, Scale bar, 10 μm. (**B**) Imaris 3D rendering of PSD-95 puncta volume in astrocyte field of views (FOVs) across brain regions. (**C**) *Left*, UPF2 deficient animals demonstrated reduced PSD-95 puncta volume across brain areas as compared to CTRL (HPC, CTRL: 1284 ± 143, 20 FOVs; cKO: 328 ± 43, 20 FOVs, p < 0.0001, Unpaired *t*-test. CTX, CTRL: 1095 ± 82, 20 FOVs; cKO: 535 ± 52, 20 FOVs, p < 0.0001, Unpaired *t*-test. PFC, CTRL: 908 ± 100, 20 FOVs; cKO: 380 ± 35, 20 FOVs, p < 0.0001, Mann-Whitney test. 4 FOVs per animal, 5 animals per genotype. *Right*, PSD-95 puncta volume summary data per animal (HPC, CTRL: 1284 ± 204, cKO: 328 ± 33, p = 0.0079, Mann-Whitney test. CTX, CTRL: 1095 ± 52, cKO: 535 ± 76, p = 0.0079, Mann-Whitney test. PFC, CTRL: 908 ± 91, cKO: 380 ± 55, p = 0.0079, Mann-Whitney test. 5 animals per genotype). (**D**) Schematic of stereotaxic delivery of AAV5-CamKIIα-EGFP virus to label hippocampal CA1 pyramidal neurons (*right image*) followed by tamoxifen dosing of CTRL and cKO animals. Representative images of dendritic spines in CTRL (*top*) and cKO (*bottom*). Scale bar, 10 μm. Astrocytic *Upf2* deficiency is associated with reduced spine density of CA1 pyramidal neurons in comparison to CTRL (*Left*, CTRL: 6.4 ± 0.18, 30 dendrites, cKO: 3.3 ± 0.15, 30 dendrites, p < 0.0001, Mann-Whitney test, 5 FOVs, 6 dendrites per animal, 5 animals per genotype). Summary data of spine density per animal (*Right*, CTRL: 6.4 ± 0.2, cKO: 3.3 ± 0.16, p = 0.0079, Mann-Whitney test, 5 animals per genotype). Data is presented as mean ± S.E.M. ** denotes p < 0.01, **** denotes p < 0.0001. **This figure demonstrates reduced PSD-95 levels associated with decreased spine density of CA1 pyramidal neurons in *Upf2* cKO brains.**

### Conditional deletion of *Upf2* in astrocytes *in vivo* impairs excitatory synaptic function

Synaptic transmission can be either enhanced, through long-term potentiation (LTP), or depressed, through long-term depression (LTD) (46). Given that dendritic spine densities are tightly linked to these long-term plasticity events (47), and we observed reductions both in spine and PSD-95 densities, we hypothesized that basal synaptic transmission and/or synaptic plasticity is altered in *Upf2* cKO conditions. We tested this hypothesis by performing electrophysiology recordings from CTRL and cKO mice. Because LTP and LTD events are well established and widely studied in hippocampus (46, 48, 49), we prepared acute hippocampal slices for CA1 field recordings (**Figure 3A**). We first assessed basal neurotransmission in the form of input-output curves to probe synaptic strength. Field excitatory postsynaptic potentials (fEPSPs) were elicited by increasing the stimulation intensity while measuring the afferent fiber volley amplitude. In doing so, input-output curves were generated to compare synaptic strength in CTRL and cKO mice. Our results demonstrated that fEPSP slope values were significantly lower in NMD-deficient animals as compared to CTRL (**Figure 3B**). We also examined presynaptic function by determining the paired-pulse ratio of fEPSP_2_/fEPSP_1_ synaptic responses. Our analysis revealed no significant differences across genotypes (**Figure 3C**).

**Figure 3.**
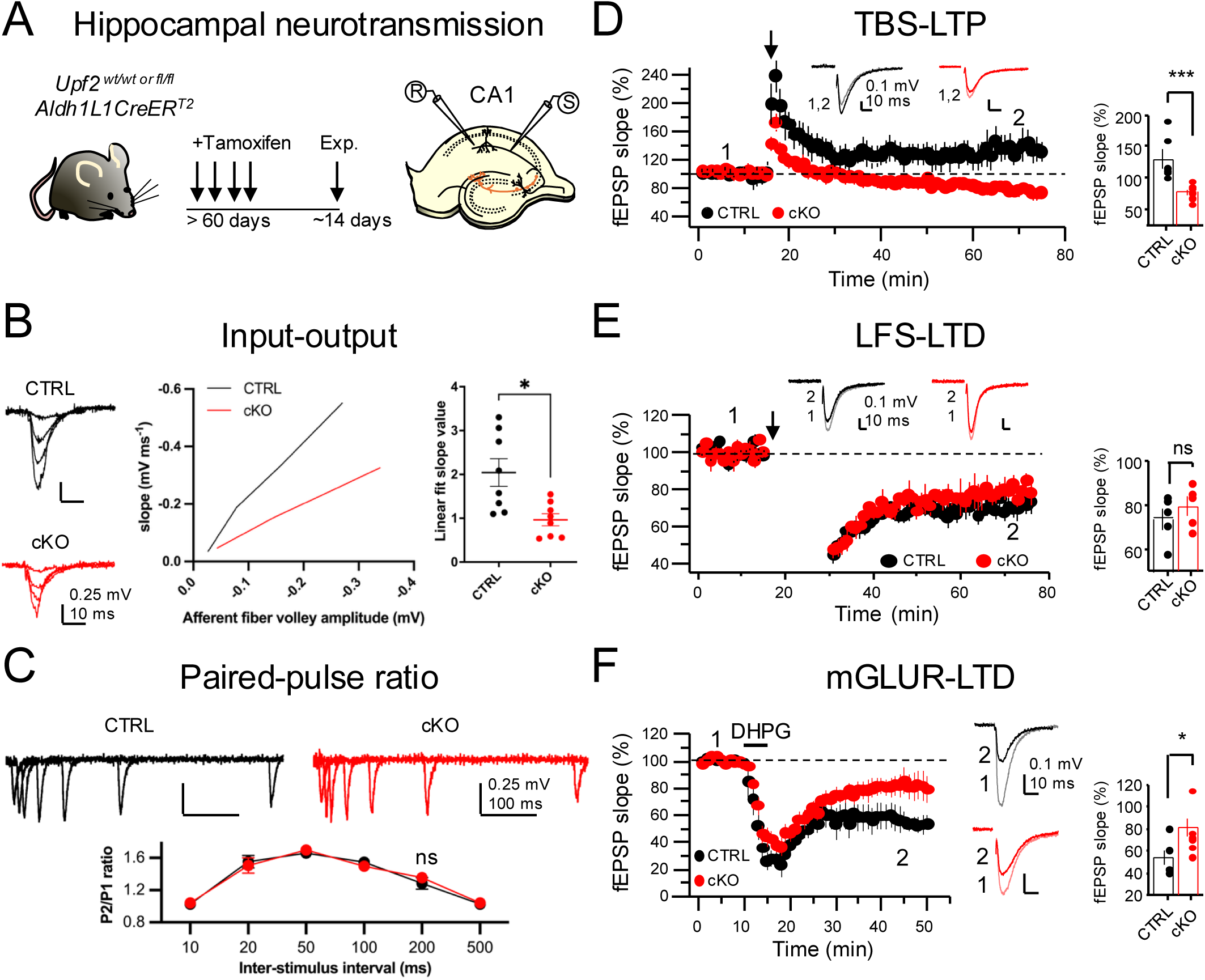
Astrocytic *Upf2* absence impairs basal neurotransmission and synaptic plasticity. (**A**) Schematic of tamoxifen dosing and electrophysiology in hippocampal area CA1 of acute brain slices. (**B**) Input-output curve assessment demonstrated reduced synaptic strength in *Upf2* cKO mice as compared to CTRL (CTRL: Linear fit 2.04 ± 0.3, r = 0.99; cKO: Linear fit 0.96 ± 0.02, r = 0.99, n = 8 slices, 4 animals per genotype, linear fit slope values: p = 0.011, Unpaired *t*-test). (**C**) Presynaptic function assessed by the paired-pulse ratio showed no differences across genotypes (CTRL, 200 ms: 1.3 ± 0.07; cKO, 200 ms: 1.35 ± 0.04, n = 8 slices, 4 animals per genotype, p = 0.4, Unpaired *t*-test). (**D**) LTP induced by theta-burst stimulation led to absent potentiation in the cKO condition (CTRL: 138 ± 15%, n = 6 mice, 12 slices; cKO: 78.8 ± 9%, n = 5 mice, 9 slices; p = 0.003, Unpaired *t*-test). (**E**) LTD induced by low-frequency stimulation was similar in both genotypes (CTRL: 72.9 ± 5%; cKO: 78.5 ± 4%, n = 5 slices, 3 animals per genotype, p = 0.41, Mann-Whitney test). (**F**) Chemical-LTD induced by DHPG (50 μM, 5 min) revealed dampened synaptic depression in *Upf2* cKO brain slices (CTRL: 55.9 ± 6%, n = 4 mice, 6 slices; cKO: 84.7 ± 9%, n = 5 mice, 7 slices, p = 0.02, Unpaired *t*-test). Data are presented as mean ± S.E.M. * denotes p < 0.05, *** denotes p < 0.005, ns = not significant. **This figure illustrates that *Upf2* deficiency in astrocytes alters synaptic function.**

Because LTP magnitude is linked to basal synaptic strength, we next assessed LTP in *Upf2* cKO hippocampus. To do this, we blocked inhibitory transmission using 100 μM picrotoxin + 3 μM CGP-55845. A 15 min baseline period of fEPSP slope values was acquired followed by LTP induction using a theta-burst stimulation (TBS) protocol. The last 10 min of fEPSP slope values were normalized to baseline to determine LTP magnitude. Our results indicated that NMD deficiency in astrocytes abolished LTP in comparison to CTRL astrocytes (**Figure 3D**). We also evaluated LTD by eliciting a low-frequency stimulation (LFS) protocol and found no significant differences in LTD magnitude in cKO slices compared to CTRL slices (**Figure 3E**).

G-protein coupled receptor (GPCR) LTD and LFS elicited LTD are distinct mechanisms for triggering long-term synaptic plasticity (50, 51). Moreover, previous studies have shown the complex impact of astrocyte signaling on GPCR-mediated LTD phenomena (51). Hence, we induced a GPCR-mediated chemical form of LTD using the Group 1 metabotropic glutamate receptor (mGluR) agonist DHPG (50 μM, 5 min). Under these experimental conditions, we observed reduced mGluR-LTD magnitude in *Upf2* cKO mice in comparison to CTRL (**Figure 3F**). Cumulatively, these results indicate that disrupted astrocytic RNA degradation leads to altered synaptic strength and attenuation of specific forms of synaptic plasticity.

### Conditional deletion of *Upf2* in astrocytes elicits anxiety-like behavior in adult mice

The prefrontal cortex, visual cortex, and hippocampus are well-characterized brain regions supporting cognitive functions that influence animal behavior (52–55). The impaired synaptic strength and plasticity in *Upf2*-cKO mice (**Figure 3**) led us to hypothesize that disrupted neural circuit activity limited cognitive processing to alter animal behavior. To test this hypothesis, we used an array of behavioral testing paradigms. Prior to initiating behavioral experiments, no difference in animal body weight was detected across test groups (**Figure S5A**). First, we implemented an Open Field Test to assess general locomotion as previously described (23, 56, 57) across genotypes (**Figure 4A**). Both groups performed similarly in this test, indicating that general motor activity is not impaired in *Upf2*-cKO mice.

**Figure 4.**
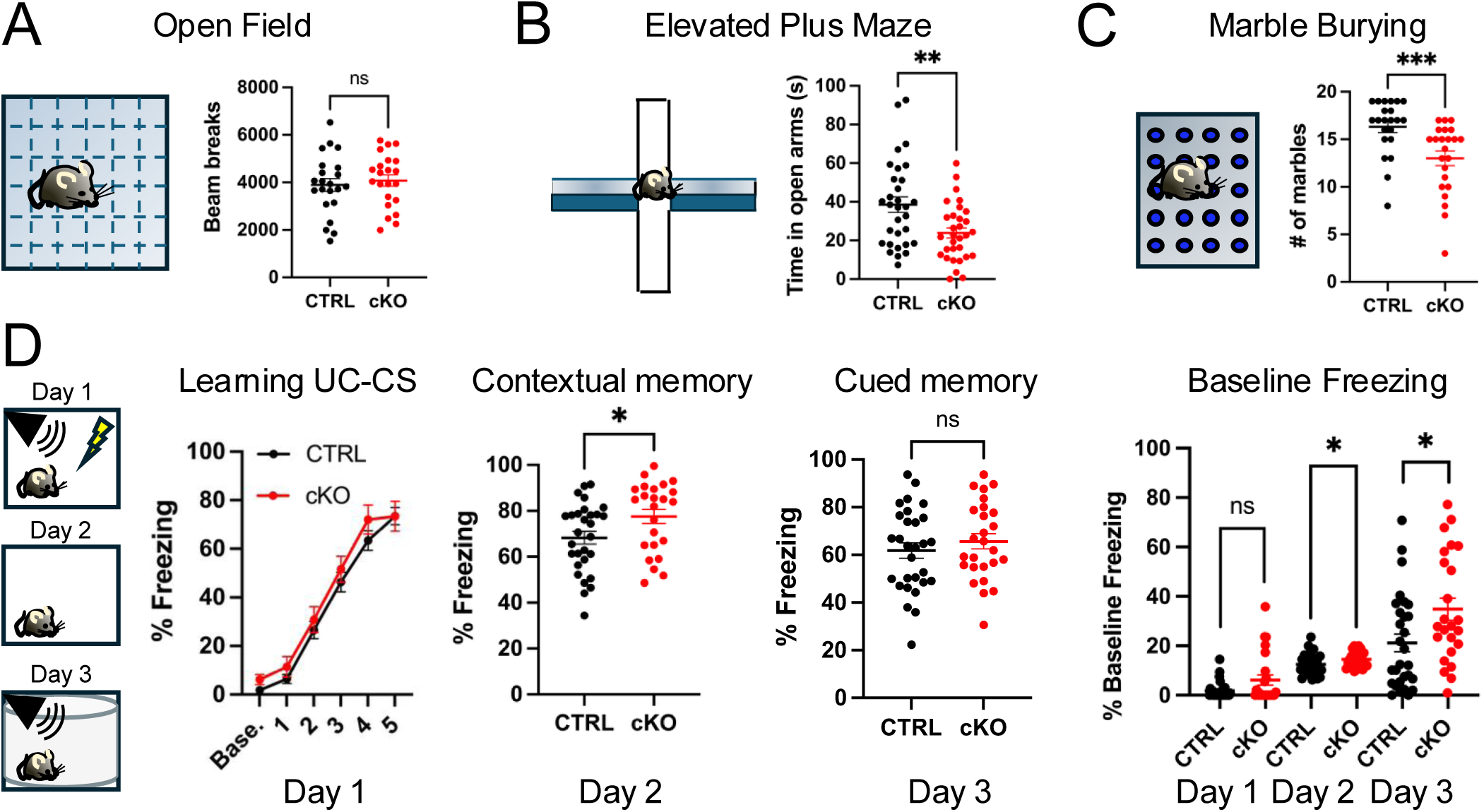
*Upf2* cKO mice display anxiety-like behavior. (**A**) Open field beam breaks display no differences in locomotion amongst CTRL and cKO animals (CTRL: 3892 ± 271; cKO: 4076 ± 239, n = 22 mice per group, p = 0.61, Unpaired *t*-test). (**B**) *Upf2* cKO mice show reduced time in open arms of elevated plus maze indicating anxiety-like behavior (CTRL: 38.5 ± 3.9; cKO: 23.09 ± 2.6, n = 31 mice per group, p = 0.0036, Unpaired *t*-test). (**C**) Astrocytic *Upf2* loss is associated with reduced marble burying (CTRL: 16.3 ± 0.6, n = 22 mice; cKO: 13.0 ± 0.8, n = 23 mice, p = 0.0003, Mann-Whitney test). (**D**) Contextual fear conditioning paradigm revealed differences in contextual memory on Day 2 between CTRL and cKO animals (CTRL: 68.3 ± 3, n = 29 mice; cKO: 77.6 ± 3, n = 24 mice, p = 0.02, Mann-Whitney test). No differences were detected on Day 1 and Day 3 (Day 1: Baseline vs. UC-CS pairings, CTRL vs. cKO, p > 0.59, Sidak’s multiple comparisons test; Day 3: CTRL vs. cKO, p = 0.69, Unpaired *t*-test). Assessment of baseline freezing revealed *Upf2* cKO mice show significant elevation on Day 2 and Day 3 as compared to CTRL (Day 1: CTRL: 1.8 ± 0.6; cKO: 6.1 ± 2.1, p = 0.43, Mann-Whitney test. Day 2: CTRL: 12.4 ± 0.8; cKO: 14.6 ± 0.7, p = 0.048, Unpaired *t*-test. Day 3: CTRL: 21.2 ± 3.6; cKO: 34.9 ± 4.5, p = 0.03, Mann-Whitney test. CTRL: n = 29 mice, cKO: n = 24 mice). Data are presented as mean ± S.E.M. * denotes p < 0.05, ** denotes p < 0.01, *** denotes p < 0.005, ns = not significant. **This figure illustrates that astrocytic UPF2 depletion is associated with avoidance behavior and increased baseline freezing.**

Astrocytes have been reported to modulate anxiety by regulating synaptic activity (58). To test whether *Upf2*-cKO mice have a defect in anxiety behavior, we used the Elevated Plus Maze (EPM) test (59). The EPM test we used contains two open arms and two closed arms, and tests mice’s natural curiosity along with their tendency to stay in enclosed spaces and avoid open spaces and heights. Anxious animals will shift the balance away from curiosity and towards safety; thus they will spend more time in the closed arms (59). *Upf2* cKO mice exhibited a strong reduction in time spent in open arms compared to CTRL mice, suggesting they exhibit anxiety-like behavior (**Figure 4B**).

Astrocytes have also been linked to regulation of repetitive behaviors (60, 61). Thus, we also conducted the marble-burying test. *Upf2* cKO mice buried significantly less marbles compared to CTRL mice (**Figure 4C**). To further probe repetitive behavior, we implemented the water spray test to induce self-grooming. Similarly, we observed a reduction in self-grooming in cKO mice (**Figure S5B**). While exhibiting the opposite of repetitive behavior, these results suggest a rather complex behavior with possible modulation of motor activity in specific environmental contexts (62).

Evidence suggests that by fine-tuning LTP, astrocytes promote the formation of new memories (63, 64). Given that LTP is impaired in cKO brains (**Figure 3**), we tested different aspects of memory. First, we probed short-term memory by implementing the Y-maze that allows an animal to explore two accessible maze arms while a designated third arm (novel arm) is blocked (**Figure S5C**). No significant differences in novel arm preference index were observed in *Upf2*-cKO mice in comparison to CTRL mice (**Figure S5C**) suggesting intact short-term memory.

To examine associative memory, we performed the 3-day contextual fear conditioning paradigm whereby an animal receives a foot shock (unconditioned stimulus, UC) paired with an auditory tone (conditioned stimulus, CS) in a designated context as previously described (23, 56, 57). On Day 1, UC-CS pairings trigger animals to freeze, the time spent freezing is quantified and interpreted as an animal’s ability to execute associative learning. Day 2 measures contextual fear memory, evaluating the animal’s freezing behavior in the same context, and Day 3 assesses cued fear memory, testing the animal’s response to the tone (CS) that was previously paired with the foot shock (UC). On Day 1, *Upf2*-cKO and CTRL mice displayed similar associative learning aptitude (**Figure 4D**). To accurately interpret freezing behavior across test days, assessing baseline freezing during the initial epoch across trials is crucial. Intriguingly, *Upf2*-cKO mice exhibited significantly higher baseline freezing responses compared to CTRL mice on Day 2-3 (**Figure 4D**), a phenotype typically interpreted as elevated baseline fear. Although we detected a significant increase in freezing behavior in *Upf2*-cKO mice during contextual memory assessment (Day 2), pre-existing baseline freezing accompanied with the anxiety phenotype (**Figure 4B**) impeded an accurate interpretation of associative memory in these mice. Higher baseline freezing has been linked to a rodent model of anxiety (65) consistent with the EPM observation for *Upf2*-cKO mice (**Figure 4B**). Together, these experiments revealed that astrocytic disruption of UPF2-dependent NMD leads to specific behavioral alterations in adult mice.

### Conditional deletion of *Upf2* in astrocytes *in vivo* upregulates mRNAs encoding signaling proteins

While our knowledge of NMD targets in neuronal populations of the central nervous system is growing, the identity and modulation of astrocytic transcripts by NMD is currently unclear (31). To gain mechanistic insight into NMD-mediated astrocyte regulation, we assessed mRNA quantities in NMD-deficient astrocytes. To do this, we isolated recombined hippocampal and cortical astrocytes using Fluorescence Activated Cell Sorting (FACS) from CTRL and cKO animals expressing ZsGreen1 (**Figure 5A**). Unbiased quantification of recombined cells during sorting determined no differences in sorted cell numbers between the groups (**Figure S6A**). Additional quantification of GFAP^+^ and S100B^+^ astrocytes in lower layers of cortex and hippocampus revealed no significant differences in cell number across genotypes (**Figure S6B**). Lastly, the canonical apoptosis marker cleaved-Caspase 3 revealed no differences amongst CTRL and cKO astrocytes (**Figure S6C**). Together, these observations strongly suggested that *Upf2* deletion did not alter cell proliferation or cell survival.

**Figure 5.**
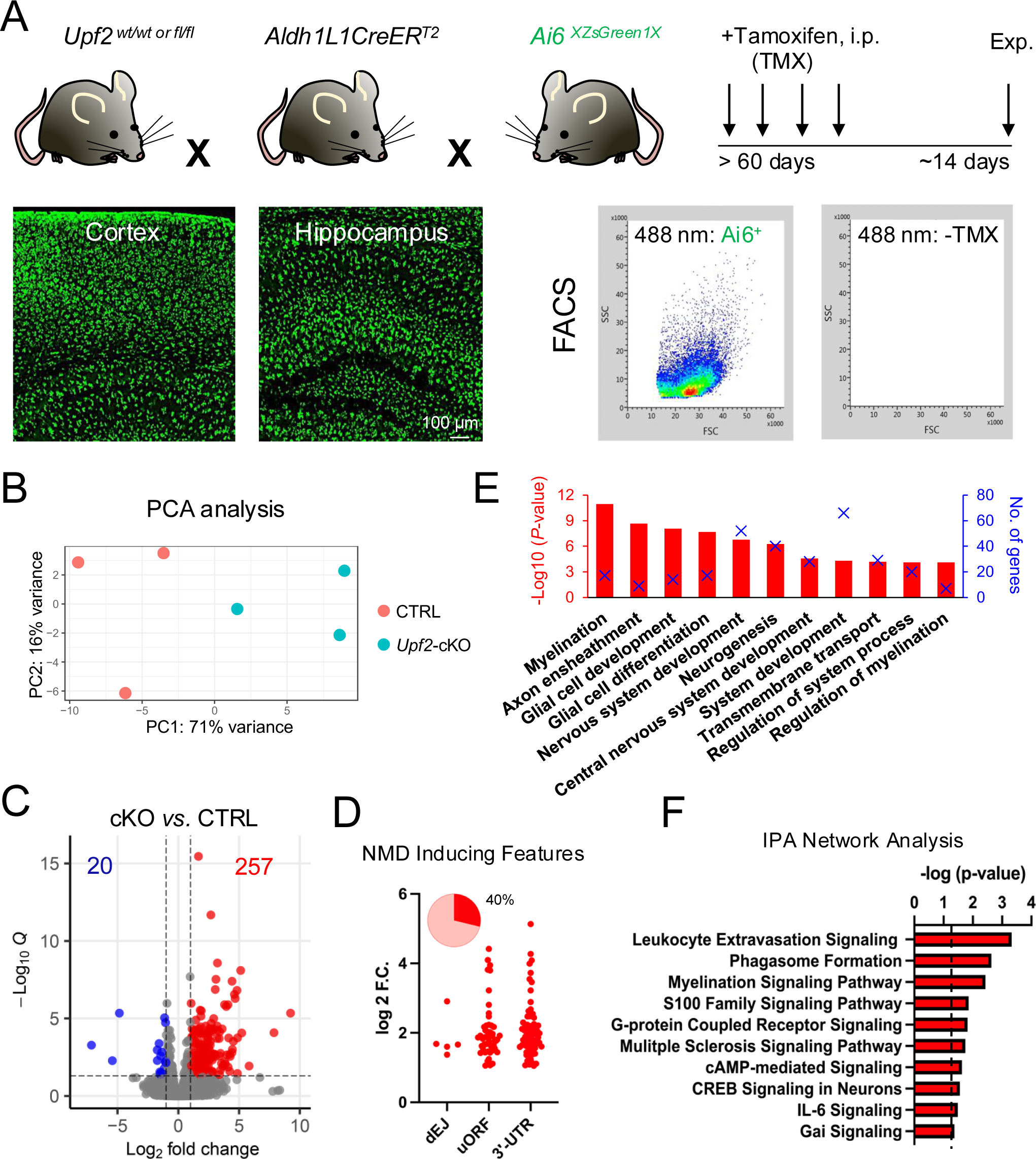
Identifying the physiological astrocytic targets of NMD *in vivo.* (**A**) *Upf2* wt/wt or fl/fl mice were crossed with the Aldh1L1Cre^ERT2^ and Ai6-ZsGreen1 transgenic lines to select CTRL and cKO astrocytes with 488 nm fluorescence after tamoxifen dosing using FACS. Representative image demonstrating ZsGreen1 expression in cortical and hippocampal astrocytes (*bottom, left*). Side scatter (SSC) and forward scatter (FSC) plot for 488 nm fluorescence selection of ZsGreen1^+^ astrocytes and tamoxifen negative (-TMX) cell suspension showing no 488 nm positivity (*bottom, right*). (**B**) Principal component analysis (PCA) of RNA-seq datasets from CTRL and cKO astrocytes, as defined in panel (A). (**C**) Differentially expressed genes (DEGs; *q* <0.05, fold change >2) identified from the RNA-seq analysis of CTRL and cKO astrocytes in panel (B). (**D**) Upregulated DEGs containing at least one NMD inducing feature: downstream Exon-Junction complex-dEJ, upstream Open Reading Frame-uORF, and long 3’-untranslated region-UTR length (>1500 nt). (**E**) Top biological functions associated with upregulated genes defined in panel (C). Statistical significance [−Log_10_ (*p*-value)] is indicated by the bar (*bottom*). The number of DEGs for a given category is indicated by an “X” (*top*). (**F**) Ingenuity Pathway Analysis (IPA) of upregulated genes defined in panel (C). Statistical significance [−Log_10_ (*p*-value)] is indicated by the bar (*top*), with a *p*-value <0.05 as the cut off. **This figure, for the first-time, reveals the physiological astrocytic transcripts regulated by NMD *in vivo* and their associated biological pathways.**

To identify mRNAs regulated by UPF2, enriched astrocyte populations from cKO and CTRL groups were subjected to RNA sequencing (RNA-seq) analysis (**Figure 5B**). This analysis identified 277 differentially expressed genes (DEGs; *q* <0.05, fold change >2) (**Figure 5C**; see supplementary data for the full list of DEGs). Because NMD facilitates mRNA degradation, its deficiency is predicted to lead to the accumulation of NMD-target mRNAs. Consistent with this notion, ∼93% of the DEGs (257 of 277) exhibited upregulation while only ∼7% (20 of 277) were downregulated in cKO astrocytes. The 257 upregulated mRNAs in cKO astrocytes are candidates to be direct NMD targets in these cells. To assess this, we screened them for 3 features mostly known to elicit NMD: an exon-exon junction downstream of the main ORF (a “dEJ”), an ORF upstream of the main ORF (an “uORF”), and a 3’UTR longer than 1,500 nucleotides (nt) (16, 66) (see Introduction). We found that 104 of the upregulated transcripts (40% of the total) contained at least one of these canonical NMD-inducing features (**Figure 5D**). Thus, these 104 mRNAs are likely NMD target mRNAs in adult cortical and hippocampal astrocytes. Many of the remaining 153 upregulated mRNAs may also be NMD targets; for example, it is known that 3’UTRs shorter than the cut-off length we selected (1,500 nt) can elicit NMD (67–72). It is also possible that some of these upregulated RNAs are indirectly regulated by NMD.

To predict biological functions regulated by NMD in adult astrocytes, we performed Gene Ontology (GO) analysis, which revealed that genes upregulated upon the loss of UPF2 are associated with “glial cell development and differentiation,” “nervous system development,” and “neurogenesis” (**Figure 5E**). We also performed pathway analysis using Ingenuity Pathway Analysis (IPA), which showed statistically significant enrichment for “leukocyte extravasation signaling,” “phagosome formation,” and “S100 Family and G-protein coupled receptor signaling” (**Figure 5F**). Intriguingly, RT-qPCR confirmed the upregulation of *S100b*, *Gabbr2*, and *Adora1* transcripts (**Figure S7**) that are linked to Ca^2+^ activity. This raised the possibility that NMD regulates astrocyte Ca^2+^ signaling, which we tested below.

### Spontaneous Ca^2+^ activity is elevated in *Upf2*-deleted astrocytes

Astrocytes are known to influence synaptic transmission and plasticity in neurons (3–5, 63) via a Ca^2+^-dependent mechanism, but how precisely this is mediated is not well understood (73–75). Given that we found Ca^2+^-dependent processes are linked with genes regulated by NMD such as *Gabbr2* and *Adora1* in astrocytes (**Figure 5F & Figure S7**), we elected to determine whether NMD has an impact on Ca^2+^ signaling in astrocytes. To test this, we measured spontaneous Ca^2+^ dynamics upon deletion of *Upf2* in astrocytes *in vivo*. For this, we induced gene deletion in hippocampus, visual cortex, and prefrontal cortex of floxed mice via injection of AAV8-GFAP-mCherry-Cre virus. Simultaneously, we delivered AAV5-GfaABC1D-cyto-GCaMP6f virus to the same brain regions to enable cell visualization (red) and measurement of Ca^2+^ transients (green) using two-photon laser scanning microscopy in acute coronal brain slices (**Figure 6A**). In the AAV5-GfaABC1D-cyto-GCaMP6f vector, the Ca^2+^ indicator *GCaMP6f* gene is driven by an astrocyte-specific promoter, ensuring its selective expression in these cells, as shown in a previous study (76). To further ensure selective live-imaging of only CRE^+^ astrocytes, the laser wavelength was tuned to 780 nm across brain areas to identify mCherry^+^ regions of interest (**Figure S8**). Time-lapse imaging enabled the acquisition of spontaneous Ca^2+^ transients reported by cyto-GCaMP6f^+^ astrocytes that facilitated the generation of heat-maps and temporal dynamic traces for Ca^2+^ fluctuations in hippocampus, visual cortex, and prefrontal cortex (**Figure 6B-D**). Measurements of peak Ca^2+^ amplitude in hippocampus, visual cortex, and prefrontal cortex revealed elevated spontaneous Ca^2+^ activity in *Upf2*-cKO slices as compared to CTRL slices (**Figure 6B-D**). These results suggest that NMD modulates spontaneous Ca^2+^ activity in mouse astrocytes and are consistent with an enrichment of Ca^2+^ signaling associated pathways in NMD-deficient astrocytes (**Figure 5F**).

**Figure 6.**
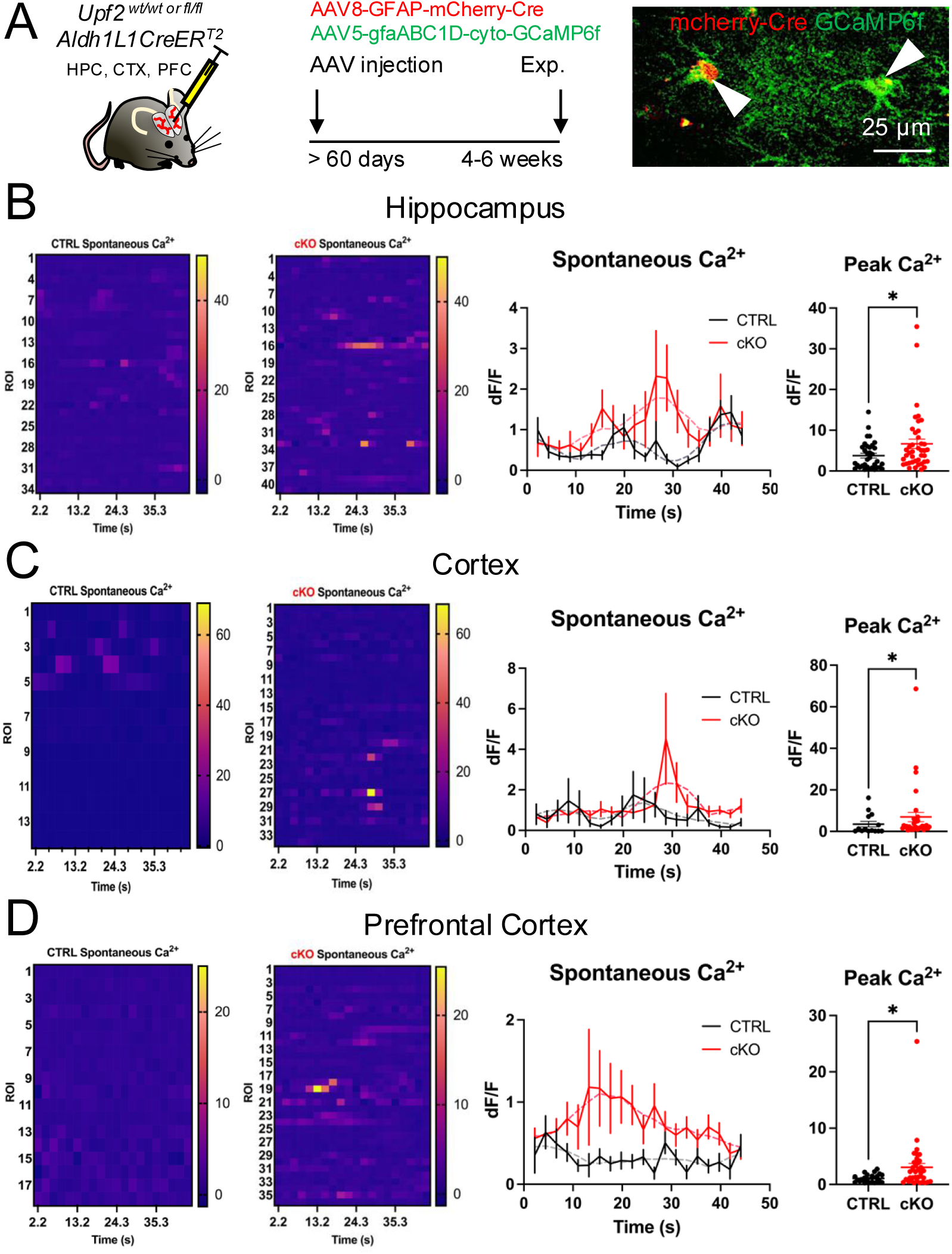
NMD*-*deficient astrocytes exhibit altered Ca^2+^ activity. (**A**) Schematic of adeno-associated viral delivery of AAV8-GFAP-mCherry-Cre/AAV5-GfaABC1D-GCaMP6f reporters to hippocampus, visual cortex, and prefrontal cortex. Representative image of mCherry/GCaMP6f expression in region of interest-ROI. In all panels of this figure: heat map of spontaneous Ca^2+^ activity in CTRL and cKO conditions (*left*), temporal Ca^2+^ dynamic traces (*middle*), and peak Ca^2+^ summary data (*right*). (**B**) *Upf2* mutant astrocytes in hippocampus show elevated peak Ca^2+^ transients of spontaneous activity in comparison to CTRL (CTRL: 3.8 ± 0.6, 34 ROIs, n = 6 mice; cKO: 6.7 ± 1.1, 41 ROIs, n = 8 mice, p = 0.038, Mann-Whitney test). (**C**) *Upf2* deficient astrocytes in visual cortex exhibit enhanced peak Ca^2+^ transients of spontaneous activity as compared to CTRL (CTRL: 3.5 ± 1.3, 14 ROIs, n = 4 mice; cKO: 6.9 ± 2.2, 34 ROIs, n = 5 mice, p = 0.046, Mann-Whitney test). (**D**) *Upf2* cKO astrocytes in prefrontal cortex display increased peak Ca^2+^ transients of spontaneous activity as compared to CTRL (CTRL: 1.1 ± 0.1, 22 ROIs, n = 4 mice; cKO: 3.0 ± 0.7, 36 ROIs, n = 5 mice, p = 0.013, Mann-Whitney test). Data are presented as mean ± S.E.M. * denotes p < 0.05. **This figure highlights that UPF2-NMD deficient astrocytes exhibit elevated Ca^2+^ transients in hippocampus, visual cortex, and prefrontal cortex.**

### Preventing high basal Ca^2+^ activity in NMD-deficient astrocytes partially restores cell volume, shifts synaptic strength, and minimizes anxiety-like behavior in *Upf2*-cKO mice

Astrocytes respond to neuronal activity by elevating their cytosolic Ca^2+^ concentrations (73, 77), which, in turn, feedback to regulate synaptic transmission strength (3–5). Consistent with this, astrocytic Ca^2+^ activity has been linked to various cognitive processes and abnormal animal behaviors (74, 78–81). This led us to hypothesize that the exaggerated Ca^2+^ activity in NMD-deficient astrocytes is responsible for altering synaptic function and thus mouse behavior. To test this, we performed a rescue experiment. We took advantage of the AAV5-GfaABC1D-hcPW2 virus carrying a Ca^2+^ extrusion (CalEx) pump shown to reduce Ca^2+^ levels in astrocytes (79, 80). We first determined whether the CalEx pump could decrease spontaneous Ca^2+^ activity in NMD*-*deficient astrocytes. Using stereotaxic surgeries, we injected CalEx, mCherry-Cre, and cyto-GCaMP6f viruses to hippocampus, visual cortex, and prefrontal cortex (**Figure 7A**). Our calcium imaging analysis revealed that the AAV5-GfaABC1D-hcPW2 virus lowered spontaneous peak Ca^2+^ transients in *Upf2-*cKO astrocytes to CTRL levels across all brain areas tested (**Figure 7B**). We next asked if reducing Ca^2+^ to CTRL levels in *Upf2-*cKO mice was sufficient to reverse defects in these NMD-deficient mice. To do this, we generated CTRL + CalEx and cKO + CalEx conditions while CTRL and cKO groups received no CalEx (**Figure 7C**). Across all test groups, we targeted viral delivery to hippocampus, visual cortex, and prefrontal cortex. To determine rescue effects, we first assessed PSD-95 levels, astrocyte cell volume, and synaptic strength. Modulation of Ca^2+^ activity in *Upf2* cKO astrocytes did not restore PSD-95 to CTRL levels while astrocyte cell volume demonstrated a partial recovery (**Figure 7D**) supporting the notion that morphological adaptations precede synaptic alterations. Nonetheless, given the upward trend of PSD-95 levels in CalEx conditions, we sought to evaluate synaptic strength. Consistent with the modest rise in PSD-95, CalEx manipulation shifted synaptic strength towards CTRL values in cKO acute hippocampal slices (**Figure 7E**). In contrast, CalEx expression in CTRL mice did not shift synaptic strength albeit a significant decrease in fiber volley amplitude (**Figure S9**). Lastly, we tested cKO + CalEx mice for the previously observed anxiety-like phenotype (**Figure 7F**). As in the previous experiments (**Figure 4B**), cKO mice exhibited low open arm time compared to CTRL mice in the EPM trial, confirming an anxiety-like phenotype (**Figure 7F**). However, cKO mice manipulated with CalEx performed similarly to CTRL mice in the EPM indicating a partial rescue of anxiety-like behavior (**Figure 7F**). Intriguingly, CTRL mice injected with CalEx displayed a reduction in open arm time suggesting an anxiogenic effect (**Figure 7F**) highlighting the complex impact of astrocyte Ca^2+^ levels on animal behavior. Altogether, these results suggest that increased Ca^2+^ activity in NMD-deficient astrocytes is, responsible for disrupting basal neurotransmission and Ca^2+^ modulation can shift synaptic strength and anxiety-like phenotypes in *Upf2* cKO mice.

**Figure 7.**
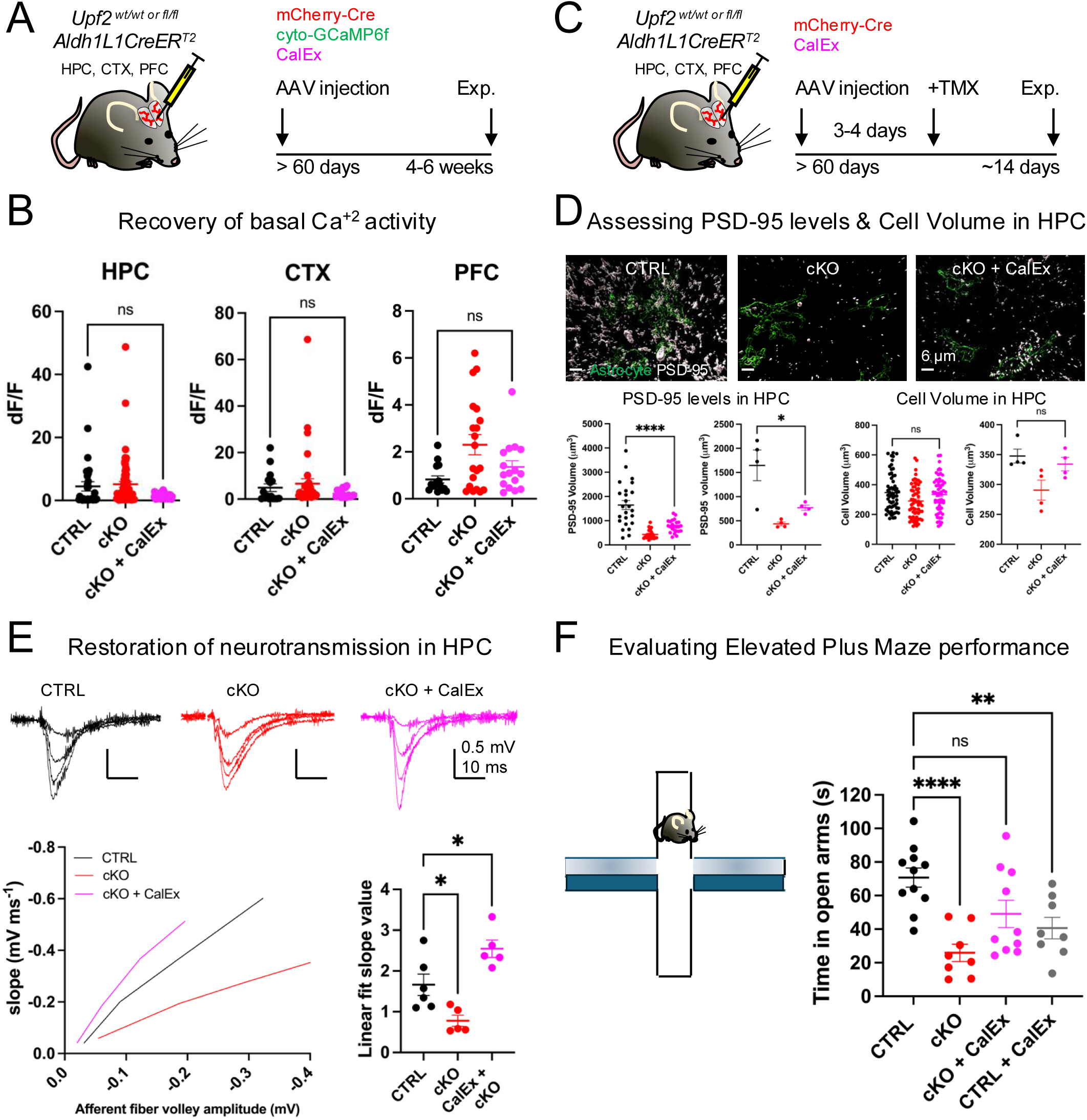
CalEx modulates basal Ca^2+^ activity in *Upf2* cKO astrocytes to partially restore cell volume, shift synaptic strength, and minimize anxiety-like behavior. (**A**) Scheme for viral delivery and timeline for Ca^2+^ measurement experiments. (**B**) CalEx diminished Ca^2+^ events to CTRL levels in hippocampus (CTRL: 4.5 ± 1.5, n = 6 mice, 32 ROIs; cKO: 5.1 ± 0.9, n = 8 mice, 68 ROIs; cKO + CalEx: 1.4 ± 0.1, n = 4 mice, 46 ROIs; One-way ANOVA DF = 2, F = 4.802, p = 0.0096, post-hoc Tukey’s multiple comparisons test CTRL vs. cKO + CalEx, p = 0.096). CalEx Ca^2+^ events are comparable to CTRL in visual cortex (CTRL: 4.9 ± 1.7, n = 4 mice, 16 ROIs; cKO: 6.6 ± 2.1, n = 6 mice, 36 ROIs; cKO + CalEx: 2.3 ± 0.4, n = 4 mice, 19 ROIs; One-way ANOVA DF = 2, F = 1.261, p = 0.29). CalEx Ca^2+^ events in prefrontal cortex are similar to CTRL (CTRL: 0.8 ± 0.2, n = 4 mice, 15 ROIs; cKO: 2.3 ± 0.4, n = 4 mice, 20 ROIs; One-way ANOVA DF = 2, F = 5.165, p = 0.0093, post-hoc Tukey’s multiple comparisons test CTRL vs. cKO + CalEx, p = 0.542). (**C**) Scheme for viral delivery and tamoxifen treatment followed by rescue assays for PSD-95 puncta volume, astrocyte cell volume, synaptic strength, and animal behavior. (**D**) *Left*, in hippocampus PSD-95 puncta volume is not restored to CTRL levels by CalEx manipulation in cKO mice (CTRL: 1648 ± 188, 24 FOVs; cKO: 437 ± 38, 24 FOVs; cKO + CalEx: 771 ± 52, 24 FOVs, 6 FOVs per animal, 4 mice per genotype; One-way ANOVA DF = 2, F = 10.28, p < 0.0001, post-hoc Tukey’s multiple comparisons test CTRL vs. cKO + CalEx, p < 0.0001). Summary data per animal (CTRL: 1648 ± 318; cKO: 437 ± 38; cKO + CalEx: 771 ± 53, 4 mice per genotype, One-way ANOVA DF = 2, F = 11.14, p = 0.0037, post-hoc Tukey’s multiple comparisons test CTRL vs. cKO + CalEx, p = 0.0223. *Right*, CalEx manipulation in *Upf2* cKO mice partially restores astrocyte cell volume assessed by S100B immunolabeling (CTRL: 348 ± 16, 60 cells; cKO: 291 ± 15, 60 cells; cKO + CalEx: 334 ± 15, 60 cells, 15 cells per animal, 4 animals per genotype, One-way ANOVA DF = 2, F = 3.75, p = 0.025, post-hoc Tukey’s multiple comparisons test CTRL vs. cKO + CalEx, p = 0.809). Summary data per animal (CTRL: 348 ± 12; cKO: 291 ± 17; cKO + CalEx: 334 ± 11, 4 animals per genotype, One-way ANOVA DF = 2, F = 4.89, p = 0.036, post-hoc Tukey’s multiple comparisons test CTRL vs. cKO + CalEx, p = 0.765). (**E**) CalEx shifts synaptic strength in hippocampus of *Upf2* cKO mice as compared to CTRL (CTRL: Linear fit 1.88 ± 0.1, r = 0.99, n = 6 slices, 3 mice; cKO: Linear fit 0.84 ± 0.05, r = 0.99, n = 5 slices, 4 mice; cKO + CalEx: Linear fit 2.66 ± 0.24, r = 0.98, n = 5 slices, 3 mice. One-way ANOVA DF = 2, F = 15.08, p = 0.0004, post-hoc Tukey’s multiple comparisons test CTRL vs. cKO + CalEx, p = 0.033). (**F**) CalEx increases elevated plus maze open arm time in *Upf2* cKO mice (CTRL: 70.7 ± 5.7, n = 11 mice; cKO: 25.9 ± 5.2, n = 8 mice; cKO + CalEx: 49.1 ± 8.1, n = 10 mice; CTRL + CalEx: 40.6 ± 6.4, n = 8 mice, One-way ANOVA DF = 3, F = 8.24, p = 0.0003, post-hoc Tukey’s multiple comparisons test CTRL vs. cKO + CalEx, p = 0.086). Data are presented as mean ± S.E.M. * denotes p < 0.05, ** denotes p < 0.01, **** denotes p < 0.0001, ns = not significant. This figure depicts that restoring Ca^2+^ levels in astrocytes of *Upf2* cKO mice partially recovers cell volume, shifts synaptic strength and minimizes anxiety-like behavior.

## DISCUSSION

Despite being well established that astrocytes perform several critical functions in the brain, the molecular mechanisms governing astrocyte activity are poorly understood. Intracellular Ca^2+^ has been widely accepted to play a vital role in encoding and transmitting information within astrocytes and thereby regulating many functions of these cells. However, molecular programs regulating astrocytic Ca^2+^ dynamics remain to be fully identified. Here, we identified the RNA degradation pathway, NMD, as a key brake that prevents abnormally high basal Ca^2+^ activity in astrocytes. Our results highlight that astrocytic RNA degradation plays a crucial role in maintaining neuro-glia interactions that support proper brain function.

RNA regulation in higher brain function has been mostly studied at the level of RNA translation in neurons. Only in the last decade neuronal RNA degradation has been recognized as a regulator of spatiotemporal gene expression to support synaptic plasticity and cognitive function. Therefore, one of the most intriguing aspects of our findings is providing evidence that astrocytic RNA degradation, similar to neuronal RNA degradation, also contributes to regulation of synaptic function and behavior via neuro-glia interactions.

Our study has revealed that NMD is required in astrocytes to sustain their size. Astrocyte’s unique morphology and interactions with synapses play a crucial role in regulating and maintaining synaptic plasticity (82, 83). Across brain structures, we observed cell volume was significantly reduced in NMD-deficient astrocytes (**Figure 1**). This is consistent with the finding that several putative NMD-target transcripts encode proteins involved in cell size regulation and/or morphology (see below).

Given that a single mouse astrocyte can interact with ∼100,000 synapses (1), alterations to astrocyte cytoarchitecture have been reported to destabilize neuro-glia interactions. Indeed, we found that PSD-95 levels of excitatory neurons are decreased across brain areas in cKO mice (**Figure 2**). This phenotype was accompanied with reduced dendritic spine density in CA1 pyramidal neurons of the hippocampus (**Figure 2**). In line with the depletion of synapse density, electrophysiological measurements revealed a decline in synaptic strength (**Figure 3**). These findings support a model that by regulating astrocytic remodeling, NMD supports synaptic architecture, leading to proper synapse density and strength. In this regard, we observed that modulation of Ca^2+^ activity in *Upf2* cKO astrocytes partially restored cell volume (**Figure 7**) supporting the notion that morphological adaptations precede synaptic alterations. A non-mutually exclusive possibility is that NMD modulates astrocytic phagocytic activity whose disruption leads to reductions in synapse density and overall strength. Temporal examination of these cellular adaptions may aid to disambiguate cooperative or specific spatiotemporal effects on synapse structure-function resulting from NMD disruption in astrocytes.

We also identified astrocytic NMD as a regulator of synaptic plasticity. *Upf2*-cKO mice demonstrated a lack of synaptic potentiation in response to physiological patterns of neural activity that trigger LTP (**Figure 3**). Recently, astrocyte remodeling of perisynaptic processes was shown to impact LTP magnitude (82), highlighting the critical role of neuro-glia structural modifications in synaptic function (83–85). Intriguingly, previous studies have shown that astrocytic S100B protein and interleukin-6 (IL-6) manipulations can downregulate LTP magnitude (86, 87) and these biological pathways were identified in our bioinformatic analysis (**Figure 5F**). Similarly, *Upf2*-cKO slices displayed reduced mGluR-LTD indicating modifications to GPCR signaling another bioinformatic biological pathway detected in our analysis (**Figure 5**). Future studies may delineate the specific crosstalk of neuro-glia GPCR signaling influencing synaptic plasticity mechanisms.

Recent evidence suggests that NMD-target mRNAs differ across cell types (ie. embryonic vs. neural progenitor cells) and may display brain-region specificity (31). Additionally, NMD target mRNAs have been mostly identified in neural stem cells and neurons in previous studies. Thus, to gain mechanistic insight into the functions of NMD in astrocytes, it is critical to identify NMD targets in these cells. We accomplished this by combining conditional deletion of *Upf2* and next-generation RNA sequencing. Consistent with the RNA degradation function of NMD, more than 90% of DEGs exhibited upregulation upon disruption of NMD in astrocytes. Indirect NMD targets, likely together with the direct targets with non-canonical-yet to be identified-NMD-inducing features, could comprise the remaining 60%. We found that NMD regulates specific mRNAs that exhibit critical functions in astrocyte cytoarchitecture and Ca^2+^ signaling in astrocytes. For example, *S100b* is implicated in astrocyte shape and migration (88). We identified *S100b* as an upregulated DEG that did not carry a canonical-NMD inducing feature and RT-qPCR confirmed *S100b* transcript upregulation (**Figure S7**). Intriguingly, the release of S100B protein can influence direct or indirect phagocytic activity by inducing proinflammatory pathways that could remodel neuro-glia interactions (89). This is consistent with upregulation of transcripts linked to S100 Family signaling pathway and IL-6 signaling in *Upf2* cKO astrocytes. Similarly, *Cldn9, Cldn11, and Mmp8* are upregulated DEGs suggested to mediate morphological adaptations in various conditions (90–92). Among them, *Cldn9* contains a uORF and has the potential to be directly targeted by NMD. Meanwhile, *Gabbr2,* encoding for GABAB, and *Adora1*, encoding for A1 receptor, stand out as GPCRs implicated in downstream Ca^2+^ signaling effector pathways that could influence neuro-glia communication. Notably, both *Gabbr2* and *Adora1* were detected as upregulated candidates containing canonical NMD-inducing features and RT-qPCR showed *Gabbr2* and *Adora1* transcript upregulation in NMD deficient conditions (**Figure S7**). Moreover, while our NMD transcript analyses identify molecular players involved in neuro-glia interactions supporting brain function, it is important to consider cell-compartment specific transcriptional regulation. Efforts towards characterizing compartment specific NMD targets in the cell body and branches are an important step to gain deeper insight into RNA glia biology.

Here, we report evidence that NMD functions to suppress basal Ca^2+^ activity in astrocytes. Our findings provide empirical evidence that it is critical that NMD suppress basal Ca^2+^ activity in astrocytes, as otherwise this leads to alterations in synaptic strength (**Figure 7E**) and promotes anxiety-like behavior in mice (**Figure 7F**). This is consistent with the known role of Ca^2+^ in astrocytes (58, 64, 74, 78–81). For example, abnormally high basal Ca^2+^ levels could lead to dysregulated release of gliotransmitters (e.g., ATP or glutamate) and other glia-derived molecules known to modulate neural excitability and synaptic transmission (4, 6, 93, 94). This would explain why we observed a decline in synaptic strength and limited synaptic plasticity in NMD-deficient *Upf2*-cKO mice. Previous studies have also suggested that astrocytic GABAB and A1 receptor signaling pathways contribute to Ca^2+^ fluctuations and modulate excitatory transmission, respectively (95–97).

Ultimately, maladaptive astrocyte Ca^2+^ influx may impact neural circuit function leading to alterations in animal behavior. We provide evidence that astrocytic NMD influences anxiety-like behavior in adulthood. In fact, astrocytes in different brain regions have been reported to suppress anxiety-like behavior (58, 78, 98–100) and astrocytic Ca^2+^ has been linked to anxiety (58). Specifically, extruding Ca^2+^ from nucleus accumbens astrocytes by overexpression of the CalEx pump led to pronounced reductions in anxiety behavior and increased exploratory behavior in mice (99). In contrast, mice with chronically activated Ca^2+^ signals in ventral CA1 exhibited enhanced anxiety behavior (100). Our study identifies astrocytic NMD as a mechanism that minimizes anxiety-like behavior by modulating basal Ca^2+^ activity in astrocytes. We have previously reported that neuronal NMD has major roles in cognitive function influencing various types of learning and different aspects of memory (23). Our current study determined that NMD in astrocytes specifically contributes to regulation of anxiety-like behavior, suggesting a complementary cell-specific role for this molecular program in rodents.

In conclusion, our findings provide key insights into constructing a comprehensive model of how cell-specific mRNA degradation impacts synaptic function and animal behavior. Our studies also have broad relevance for understanding the significance of astrocytic mechanisms in synaptic plasticity, disruption of which contributes to several brain ailments. Here, we specifically disturbed NMD in astrocytes, which allowed us to evaluate the physiological roles of this highly conserved and selective RNA turnover pathway in astrocyte biology and synaptic function. However, astrocytes exhibit regional and functional heterogeneity (101, 102) and NMD may display cell-specific RNA regulatory roles driven by specific machinery proteins. While our work establishes the physiological requirement for astrocytic NMD in three critical brain regions, future studies are required to define the spatiotemporal impact of NMD on astrocyte-neuron function in the nervous system.

## MATERIALS AND METHODS

### Transgenic Animals and Tamoxifen Dosing

Animal handling followed an approved protocol by the Institutional Animal Care and Use Committees of Weill Cornell Medicine and the Albert Einstein College of Medicine in accordance with NIH guidelines. The *Upf2* conditional knockout mouse line (41) was backcrossed for at least eight generations with C57BL/6J (Stock#: 000664, The Jackson Laboratory) mice to generate *Upf2*^wt/wt^ and *Upf2*^fl/fl^ animals that were bred with *Aldh1L1CreER^T2^* (B6N.FVB-Tg(Aldh1l1-Cre/ERT2)1Khakh/J Stock#: 031008, The Jackson Laboratory (42)) transgenic mice to enable CRE expression in astrocytes of *Upf2*^wt/wt^ and *Upf2*^fl/fl^ mice in a tamoxifen dependent manner. Mice were housed in 12:12 light-dark conditions with *ad libitum* feed diet. At 2-4 months of age *Upf2*^wt/wt^:*Aldh1L1CreER^T2^*(CTRL) and *Upf2*^fl/fl^:*Aldh1L1CreER^T2^* (cKO) mice received intraperitoneal tamoxifen (T5648, Sigma-Aldrich, USA) in corn oil at 200 mg/kg for 4 consecutive days using a 26 gauge Sub-Q syringe (309597, Becton Dickinson & Co., USA). For fluorescence expression of the ZsGreen1 reporter, the Ai6 transgenic mouse line was implemented (B6.Cg-Gt(ROSA)26Sortm6(CAG-ZsGreen1)Hze/K, Stock#: 007906, The Jackson Laboratory (103)). *Upf2*^wt/wt^:*Aldh1L1CreER^T2^:Ai6* (CTRL) and *Upf2*^fl/fl^:*Aldh1L1CreER^T2^:Ai6* (cKO) followed the previously mentioned tamoxifen regime. Experimental manipulations were initiated 12-14 days after the last day of tamoxifen dosing, a timeline demonstrated to sufficiently deplete UPF2 (23, 35). Male and female mice were included for all experimental manipulations.

### Perfusion and Cryo-sectioning

Mice were anesthetized using 4% isoflurane (11695-6777-2, Covetrus) and intracardially perfused with 10 mL of 1X phosphate buffer saline-PBS (MRGF-6235-020Q, GrowCells) to flush blood, followed by ∼15 mL of 4% paraformaldehyde-PFA (15713-S, Electron Microscopy Sciences) for fixation. Following PFA perfusion, mouse brains were extracted and stored in 4% PFA at 4°C overnight. Next day, brains were rinsed with PBS and stored at 4°C in 30% sucrose (S0389, Sigma Aldrich). After achieving sucrose penetration for cryoprotection, brains were embedded in tissue trays containing OCT compound (4583, Sakara), frozen on dry ice, and stored at -80 °C until cryosectioning. A Leica CM3050S cryostat enabled the collection of frozen sections (25-70 μm thickness) mounted onto microscope slides and stored at -20 °C for downstream application.

### Immunohistochemistry and Confocal microscopy

Frozen sections were rehydrated using PBS and subsequently rinsed 3 x for 5 min on a rotating tray, 45 rpm. Tissue slides were then permeabilized and blocked with a PBS solution containing 0.2-0.5% Triton-X (X100, Sigma Aldrich) and 10% normal goat serum (G9023, Sigma Aldrich) for 1 h. After permeabilization and blocking, tissue slides were incubated at 4°C overnight with primary antibodies at specified dilutions, unless stated otherwise. Next day, tissue slides were washed with PBS 3 x for 15 min on a rotating tray, 45 rpm. Secondary antibodies were added and incubated at room temperature for 1-2 h, at specified dilutions, unless stated otherwise. In some cases, tissue slides were rinsed with PBS and incubated with Hoechst 33342 (H3570, ThermoFisher Sci) for 5 min. After 3 x 10 min PBS washes, tissue slides were allowed to air dry protected from light. ProLong Anti-Fade Glass Mountant (P36980, ThermoFisher Sci) was added, coverslips adhered, and slides were stored at 4°C. When recommended, antigen retrieval was performed by incubating tissue slides in citrate buffer, pH = 6 (C9999, Sigma Aldrich) and heated until solution boiled. The following primary and secondary antibodies were used: rabbit anti-hUPF2 1:500 (Jens Lykke-Andersen Lab), chicken anti-GFAP 1:5000 (PA1-10004, ThermoFisher Sci), chicken anti-Iba1 1:500 (234009, Synaptic Systems), rat anti-Mbp 1:1000 (ab7349, 1 h primary incubation, Abcam), mouse anit-PSD-95 1:500 (MA1-046, ThermoFisher Sci), rat anti-LAMP2 1:300 (ab13524, Abcam), guinea pig anti-S100B 1:200 (287004, Synaptic Systems), chicken anti-GFP 1:100 (A10262, ThermoFisher Sci), mouse anti-GFP 1:50 (A11120, ThermoFisher Sci), rat anti-mCherry 1:50 (M11217, ThermoFisher Sci), mouse anti-NEUN 1:500 antigen retrieval (MAB377, Sigma Aldrich), rabbit anti-ASPA 1:100 (ab223269, Abcam), rabbit anti-cleaved Caspase 3 1:500 (9664, Cell Signaling Tech), anti-rabbit Alexa Fluor 488 1:500 (A32731, ThermoFisher Sci), anti-chicken Alexa Fluor 647 1:1000-GFAP, 1:500-Iba1 (A21449, ThermoFisher Sci), anti-rat Alexa Fluor 546 1:500 (A11081, 1 h secondary incubation, ThermoFisher Sci), anti-mouse Alexa Fluor 546 1:500 (A11003, ThermoFisher Sci), anti-rat Alexa Fluor 633 1:500 (A2104, ThermoFisher Sci), anti-guinea pig Alexa Fluor 546 1:500 (A11074, Thermo Fisher Sci), anti-chicken Alexa Fluor 488 1:500 (A11039, ThermoFisher Sci), anti-mouse Alexa Fluor 488 1:500 (A21131, Thermo FisherSci), anti-rat FITC 1:500 (11-4817-82, ThermoFisher Sci), Hoechst 334222 1:5000 (H3570, ThermoFisher Sci). An Olympus Fluoview FV1000 confocal microscope enabled the acquisition of fluorescence images at 512 x 512 or 640 x 640 pixel resolution assisted with Olympus Fluoview 4.2 software. The spinning-disk confocal Andor DragonFly microscope using 1024 x 1024 pixel resolution operated with Fusion software also assisted image acquisition. Laser channel detections were set to 405 nm, 488 nm, 546 nm, and 647 nm for image acquisition using 10X (dry), or 60X (1.4 NA) oil-immersion objective with manufacturer oil (Olympus Immoil F30CC, Type-F). Images were processed in Fiji of at least four field of views per genotype.

### Fluorescence Assisted Cell Sorting

To isolate 488 nm^+^ astrocytes, *Upf2*^wt/wt^:*Aldh1L1CreER^T2^*and *Upf2*^fl/fl^:*Aldh1L1CreER^T2^* mice were crossed with the Ai6 reporter line from The Jackson Laboratory to generate triple transgenic mice. Fluorescence Assisted Cell Sorting (FACS) was performed ∼14 days after the last day of tamoxifen dosing. Mice were anesthetized with 4% isoflurane followed by intracardial perfusion using Hanks’ Balanced Salt Solution (88284, ThermoFisher Sci) to flush blood. Next, brains were extracted onto ice-cold petri dishes for cortex-hippocampus dissections. Cortical-hippocampal samples from each genotype were collected in ice-cold petri dishes containing DNase, RNase, protease free PBS (BP24384, Fisher Sci). After tissue collection (n = 3 brains/genotype), samples were manually chopped using a sterile scalpel. Cell dissociation was performed using a Papain Dissociation Kit (LK003150, Worthington Biochemical Corp). During the papain dissociation step, RNAse murine inhibitor (M0314S, New England Biolabs) was added at 1U/μL working concentration. Digested cell suspensions were passed through a 40 μm nylon mesh cell strainer (431750, Corning) followed by cell centrifugation at 300 g for 5 min. Cell pellets were gently resuspended and layered onto an ovomucoid gradient for final cell pelleting at 70 g for 6 min. CTRL and CKO cell pellets were re-suspend in 0.5% BSA (0332, VWR Chemicals) made in PBS solution for myelin removal using a magnetic bead kit (130-096-733, Miltenyi Biotec) with LS columns (130-042-401, Miltenyi Biotec). After myelin depletion, CTRL and cKO cell collections were brough to a cell sorter. A BD Influx system enabled FACS and an Ai6 negative cell suspension served as a negative control. Forward and side scattering gates were used to remove clumps of cells and debris. On average a total of 1-1.6 x 10^6^ purified cells per genotype were collected in 0.5% BSA-PBS containing RNAse murine inhibitor for downstream RNA extraction.

### RNA Sequencing and Analysis

CTRL and cKO cell samples purified by FACS from 6 mice per genotype (with 2 technical replicates) were immediately processed for RNA extraction using the RNeasy Mini plus kit (74136, Qiagen). After following the manufacturers specifications, RNA quantity and quality was assessed using Agilent Bioanalyzer Tape station and picogel analyses. RNA integrity number (RIN) values ranging from 7.0 – 9.0 for CTRL and cKO RNA samples were used to prepare the library using the IlluminaNextSeq500 SMART-Nextera kit, as per manufacturer’s instructions. Libraries were sequenced (pair-end reads) with an Illumina NextSeq 500 platform for 50 cycles, with a depth of 40M reads per sample.

As performed previously (22, 104), reads were filtered for quality and aligned with STAR (2.5.2b) against the GRCm39 (Ensembl version 105). The exon counts were aggregated for each gene to build a read count table using SubRead function featureCounts. Differentially expressed genes were defined using the DESeq2 program with the following thresholds: |Log2FC| >1 and *q*-val <0.02. The database for annotation, visualization and integrated discovery (DAVID) v6.8 was used for GO analysis. For pathway analysis, the upregulated genes, along with their fold-change and adjusted *p*-values were uploaded to Ingenuity Pathway Analysis (Qiagen) and analyzed. A *p*-value <0.05 was used as the significance threshold, and the enriched signaling pathways were categorized based on z-score values to generate predicted biological pathways.

To identify NMD-inducing features among upregulated transcripts, only Ensembl transcripts with detectable 5’UTRs and 3’UTRs were considered. Transcripts were classified as having a dEJC if they contained at least one exon-exon junction ≥ 50 nucleotides downstream of the stop codon defining the main ORF. Transcripts were considered to have an “upstream (u) ORF” if it included an ORF upstream of the main ORF that was ≥ 30 nucleotides in length and initiated by a codon in a context with a purine at the −3 position or a guanine at the +1 position (relative to the A in the AUG initiation codon [+1]). To minimize the identification of uORFs capable of reinitiating translation (and thereby escaping NMD), we required that the uORF not overlap with any sequence in the main ORF.

### RT-qPCR

From CTRL and *Upf2*-cKO FACS selected ZsGreen1^+^ astrocytes total RNA was extracted using an RNA extraction kit (R2035, Zymo) as specified by manufacturer. 200 ng of RNA from 4 cell selections from each group (n = 3-4 brains per genotype group) was transcribed into cDNA using the SuperScript VILO cDNA synthesis kit (11754050, ThermoFisher Sci). Real Time-quantitative PCR (RT-qPCR) reactions were prepared using the PerfeCTa SYBR method kit (95074-05K, Quantabio). For each reaction 10 ng of cDNA was used. Primer list: *Upf2* (Fw: AGCTGCAGAACACTGAGAGG; Rev: CTCCTCATATTGCTTGTGCC), *S100b* (Fw: AAAGGCTCATGGGCTCGAAG; Rev: GAAGGGGGTTGGGGTTTCAT), *Gabbr2* (Fw: GTTGTGCCTTTGAGGAGAGC; Rev: AGTCCACTCCGATGTAGCCT), *Adora1* (Fw: GATCGGTACCTCCGAGTCAAGA; Rev: CACTCAGGTTGTTCCAGCCAAAC), *β-actin* (Fw: CTCTTCCAGCCTTCCTTCCT; Rev: TGCTAGGGCTGTGATCTCCT). Primers were custom orders from Integrated DNA Technologies. Primer efficiencies were determined by cDNA sample dilution to achieve linear regression Ct values of Ct = 20-30. The comparative ΔΔCT method (105) was executed using a QuantStudio 3 instrument (appliedbiosystems, ThermoFisher Sci) to quantify mRNA levels. Relative expression levels were normalized to housekeeping gene *β-actin* mRNA.

### Astrocyte Imaris 3D reconstructions, PSD-95 measurements, branch length analysis

For Lck-GFP astrocyte morphology, 70 μm-thick frozen brain sections from both genotypes were immunostained with mouse anti-GFP 1:50 (A11120, ThermoFisher Sci) and rat anti-mCherry 1:50 (M11217, ThermoFisher Sci) antibodies. To quantify excitatory synapse numbers, 50 μm-thick frozen brain sections from both genotypes were immunostained with mouse anit-PSD-95 1:500 (MA1-046, ThermoFisher Sci) antibody. Fluorescence imaging was carried out on an inverted Leica DMi8 microscope equipped with Andor Dragonfly 200 unit (405 nm, 488 nm, 561 nm, 640 nm and 730 nm lasers) operated by Fusion software. For each mouse, a Z-stack field of view for each brain region (HPC, CTX, and PFC) was acquired (30 μm depth scan size, 0.2 μm interval Z-stack, with a 63X objective and 1024 x 1024 pixel resolution). Imaris 10.2.0 software facilitated 3D-surface rendering using the mask function for quantification of cell volume and PSD-95 puncta volume in blind fashion as previously described (106, 107). Inclusion criteria for astrocyte cells required nuclei to be located at least 15 μm below the surface of the Z-stack imaging plane. Astrocyte cells residing in the outer edges of the field of view were not included. For cell volume analysis, 20 cells from each mouse per genotype were collected for each brain area. To quantify PSD-95 levels, total PSD-95 puncta volume (outside and inside astrocytes) for each field of view was summed. Four field of views were collected for each mouse and each brain area for a total of 20 field of views per genotype. For CalEx PSD-95 rescue experiments in the hippocampus, 6 fields of view were acquired for each mouse for a total of 24 field of views per genotype. For CalEx cell volume rescue experiments in the hippocampus, astrocytes were labeled using the guinea pig anti-S100B 1:200 (287004, Synaptic Systems) antibody,15 cells from each mouse per genotype was acquired. For astrocyte branch length analysis, ZsGreen1^+^ astrocyte images from CTRL and cKO mice were converted to 8-bit images followed by threshold adjustment using Fiji software. Next, images were processed with the make binary function and skeletonize feature. Lastly, images were analyzed using the Skeleton-Analyze Skeleton (2D/3D) function and maximum branch length values were exported. Data was filtered by only including values greater than 2 μm and averaging the values for each field of view. For each genotype 4 field of views per animal was analyzed for cortex and hippocampus.

### Stereotaxic Surgeries

Transgenic mice were anesthetized using 4% isoflurane mixed with O_2_ flowing at 2 mL/min. Mice were placed in a Kopf stereotaxic instrument and anesthesia was sustained at 1.5-2% isoflurane-O_2_ mix. For spine density experiments, 100-200 nL of AAV5-CamKIIα-EGFP (50469, Addgene, titer: 1.9 x 10^13^) was delivered bilaterally using a Hamilton syringe at stereotaxic coordinates targeting the CA1 area of the hippocampus (A/P: -2.0, M/L: 1.5, D/V: -1.1 mm). Animals were allowed to recover from surgery for three days and then received tamoxifen administration. For astrocyte morphology experiments, 300-500 nL of AAV8-GFAP-mCherry-Cre (UNC Vector Core, titer: 4.1 x 10^12^) and AAV5-GfaABC1D.PI.Lck-GFP.SV40 (105598, Addgene, titer: 2.3 x 10^13^) mixed at a (1:2 ratio) was delivered at stereotaxic coordinates targeting prefrontal cortex (A/P: +1.8, M/L: 0.3, D/V: -1.6 mm), the CA1 area of the hippocampus (A/P: -2.0, M/L: 1.5, D/V: -1.2 mm) and visual cortex (A/P: -2.0, M/L: 1.5, D/V: - 0.75 mm). After recovering from surgery, (3-4 days) animals underwent tamoxifen administration. For astrocyte calcium imaging experiments, 1.2 μL of AAV8-GFAP-mCherry-Cre (UNC Vector Core) and AAV5-gfaABC1D-cyto-GCaMP6f (52925, Addgene, titer: 2.5 x 10^13^) mixed at a (1:2 ratio) was delivered to prefrontal cortex, hippocampus, and visual cortex using the previously mentioned stereotaxic coordinates. Two-photon imaging experiments were performed 4-6 weeks post-surgery. For CalEx calcium imaging 1.5 μL of AAV5-GfaABC1D-mcherry-hPMCA2w/b (111568, Addgene, titer: 1.2 x 10^13^) virus together with AAV5-gfABC1D-cyto-GCAMP and AAV8-GFAP-mCherry-Cre mixed at a (3:2:1 ratio) was injected at the stereotaxic coordinates previously described and experiments were performed 4-6 weeks post-surgery. For CalEx behavioral rescue experiments 1.5 μL of AAV5-GfaABC1D-mcherry-hPMCA2w/b and AAV8-GFAP-mCherry-Cre mixed at a (2:1 ratio) was delivered to prefrontal cortex, hippocampus, and visual cortex. Animals recovered from surgery for 3-4 days and underwent tamoxifen administration. All mice received meloxicam and bupivacaine following the IACUC recommendations of Weill Cornell Medicine. The EGFP, GCaMP6f and CalEx viruses were acquired from Addgene, the mCherry-Cre virus was ordered from the UNC Vector Core, Chapel Hill.

### Spine Density Assessment

Transgenic mice were monitored for 72 h post-stereotaxic surgery and then dosed with tamoxifen. After ∼14 days of the last tamoxifen dose, mice were anesthetized with 4% isoflurane and intracardially perfused with PBS to flush blood followed by 4% PFA. Brains were fixed in 4% PFA overnight and then cryoprotected with 30% sucrose. Upon achieving cryoprotection, brains were embedded in tissue trays, frozen in OCT compound, and stored at -80°C. Coronal hippocampal sections of 250-300 μm thickness were mounted onto microscope slides using a Leica CM3050S cryostat and stored at -20°C. Next day, coronal sections underwent immunostaining procedures to enhance EGFP signal with an anti-GFP chicken antibody 1:50 (A10262, ThermoFisher Sci) or mouse anti-GFP 1:50 (A11120, ThermoFisher Sci). In some cases, tissue slides were incubated with Hoechst 33342 1:5000 (H3570, ThermoFisher Sci) for 5 min then washed with PBS 3 X 5 min, to visualize brain cells. After following the immunostaining procedures previously described, tissue slides were allowed to air dry protected from light and Pro-Long Antifade Glass Mountant was added for coverslip adhesion. The Olympus Fluoview FV1000 confocal microscope enabled the acquisition of EGFP signals at 640 x 640 pixel resolution using the 488 nm laser wavelength path with a 60X (1.4 NA, Nikon) oil-immersion objective at 4.0 μs/pixel, digital zoom 8X. Z-stack images of 10-15 μm thickness with 0.2 μm steps were acquired from dendrites of hippocampal CA1 pyramidal neurons at least 150 μm away from the cell body. Images were processed in Fiji for Z-projections of at least 5 field of views visualizing 6 dendrites per animal for each genotype. Spine structures were quantified in 10 μm bins per dendritic image to determine spine density in CTRL and cKO conditions in a blinded manner.

### Hippocampal slice preparation

Male and female mice were anesthetized with 4% isoflurane followed by perfusion with 25 mL of cold NMDG solution containing in (mM): 93 NMDG, 2.5 KCl, 1.25 NaH_2_PO_4_, 30 NaHCO_3_, 20 HEPES, 25 glucose, 5 sodium ascorbate, 2 Thiourea, 3 sodium pyruvate, 10 MgCl_2_, 0.5 CaCl_2_, brought to pH 7.35 with HCl (SA49, Fisher Sci). Following perfusion, the brain was extracted and dissected hippocampi were cut using a VT1200s (Leica Co.) microslicer in cold NMDG solution. Acute hippocampal slices (300-400 μm) were collected and placed in a chamber containing extracellular artificial cerebrospinal fluid (ACSF) recording solution containing (in mM): 124 NaCl, 2.5 KCl, 26 NaHCO_3_, 1 NaH_2_PO_4_, 2.5 CaCl_2_, 1.3 MgSO_4_ and 10 glucose. The slice chamber was warmed in a water-bath at 33-34°C. After 10 min of completing slice collection, the chamber was moved to room temperature and slices were allowed to recover for at least 45 min prior to experimentation. All solutions were equilibrated with 95% O_2_ and 5% CO_2_ (pH 7.4).

### Electrophysiology

Electrophysiological experiments were performed at 31.2 ± 1 °C under temperature control (TC-344C, Warner Instruments) in a submersion-type recording chamber perfused at 2 mL/min with ACSF. A stimulating glass electrode was filled with ACSF and placed in *stratum radiatum* to activate Schaffer collateral inputs using a Isoflex stimulus isolator (A.M.P.I) with a 100 μs pulse width duration. Extracellular field excitatory postsynaptic potentials (fEPSPs) were recorded using a patch-type pipette filled with 1 M NaCl. For long-term potentiation (LTP) recordings, slices of 400 μm thickness were stimulated every 20 s for a baseline fEPSP period of 15 min. LTP was induced by theta-burst stimulation (TBS) consisting of: 10 bursts of 5 pulses at 100 Hz delivered every 200 ms (inter-burst interval) repeated 4X (every 5 s) as previously described (106). LTP experiments were carried out in the presence of picrotoxin (100 μM, P1675, Sigma Aldrich) and CGP-55845 (3 μM, No. 1248, Tocris Bioscience) to block inhibitory transmission. The last 10 min of fEPSP responses were compared to baseline average responses to determine the magnitude of potentiation. For extracellular field input-output and paired-pulse ratio experiments synaptic inputs were stimulated every 10 s. The stimulation intensity of an Isoflex stimulator was increased from 0-20 V in 5 V increments. Slopes of fEPSP responses and amplitudes of fiber volleys were determined using ClampFit 11.2 software. Paired-pulse ratio (PPR) was examined by delivering two stimuli at various inter-stimulus intervals (10-500 ms) and measuring the ratio of fEPSP slopes (fEPSP_slope2_/fEPSP_slope1_). Both input-output and paired-pulse ratio was performed in the absence of pharmacological drugs. Long-term depression (LTD) was induced using a low-frequency stimulation protocol (LFS: 900 pulses at 1 Hz). A 15 min baseline period was recorded prior to LTD induction and synaptic inputs were stimulated every 20 s. For chemical LTD, mGluR-LTD was elicited by bath-application of DHPG (50 μM, 5 min, No. 0342, Tocris Bioscience) after obtaining a 10 min baseline period. To determine the magnitude of depression for LFS and mGluR-LTD the last 10 min of fEPSP slope responses were compared to baseline averaged responses. Field recordings were registered with a MultiClamp 700B amplifier and AxonDigita 1550B (Molecular Devices) and signals were filtered at 2 kHz and digitized at 5 kHz. Stimulation and acquisition were controlled with custom software (Igor Pro 6 or Clampex 11.2).

### Two-photon laser scanning microscopy for astrocyte calcium imaging

Astrocyte calcium imaging experiments were conducted 4-6 weeks after stereotaxic surgery. An In vitro Ultima 2P microscope (Bruker Corp) with an Insight Deep See laser tuned to 780 nm was used to visualize mCherry-Cre^+^ signals at 512 x 512 pixel resolution across targeted brain areas with 6-8 mW laser power measured at the 60X objective (Nikon, 1.0 NA). Upon identification of mCherry-Cre^+^ regions of interest (ROI) the laser wavelength was tuned to 945 nm, 3-4 mW laser power, for detection of GCaMP6f signals. T-series (time-series) software (PrairieView 5.4, Burker Corp) captured spontaneous Ca^2+^ transients. Images from the time series were Z-projected to identify ROIs with standard deviations denoting Ca^2+^ fluctuations. The ROIs were analyzed in a semi-automated manner using Fiji. The average value of 3 frames was considered baseline fluorescence (F_o_), and the corresponding change in fluorescence (dF = F-F_o_/F_o_) was measured. Across ROIs, the highest dF/F at a given time point was determined as the peak Ca^2+^ amplitude analyzed in blind fashion.

### Behavioral Paradigms

For all behavior experiments, animals were allowed to acclimate to room conditions for 1 h prior to initiating assays. Behavior rooms were monitored to maintain room temperature and humidity. Experimentation was conducted in 75% white light and 25% red light. Elevated plus maze was performed in 100% white light. Equipment underwent clidox sterilization between trials and 70% ethanol was used to clean contextual fear conditioning equipment. Male mice were tested prior to female mice. *<u>Open Field</u>*: The open field assay was conducted using the SD instruments Open Field Equipment and software. A 16 x 16 beam array detected animal movement in X and Y coordinates. SD software calculated beam breaks for 12 interval phases of 5 min duration for a total trial period of 1 h. Total beam breaks was calculated by the sum of X and Y coordinates in 1 h. *<u>Marble Burying</u>*: Standard home cages 27.5 cm (L) x 19.5 cm (W) x 15 cm (H) were filled with aspen chip bedding measuring a height of ∼2 cm. Marbles arranged in a 5 x 4 (row x column) equidistant matrix assayed digging behavior. Each burying trial was 30 min in duration. Marbles covered by > 75% of aspen chip bedding met inclusion criteria for buried marbles. *<u>Elevated Plus Maze</u>*: The Elevated plus maze apparatus was positioned and set for Noldus Ethovision XT software acquisition. Animals were tracked for an exploration trial period of 5 min. Open arm, center, and closed arm durations were calculated by Ethovision software. Our Plus Maze (San Diego Instruments) contains two open arms and two closed arms and relies upon the animal’s natural tendency to stay in enclosed spaces and avoid open spaces and heights. Anxious animals will spend more time in the closed arms than less anxious animals, which will explore the open arms for a longer duration. EthoVision XT detects the center point, tail base, and nose point of the animal, allowing for accurate measurement of position and enabling us to discriminate between the subject only poking its nose around the corner or moving its entire body into one of the open arms of an elevated plus maze. *<u>Contextual Fear Conditioning</u>*: Fear Conditioning chambers (Med Associates Inc.) comprise standard grid-floored chambers that are enclosed within a sound-attenuated cubicle. Each cubicle has its own lux-control box, standalone aversive stimulator, speaker (for conditioned-stimulus exposure) and high-speed firewire infrared video camera. The context of chambers can thus be modified to differ in lux, bedding (stored under grid-flooring), shape (by means of a Med Associates supplied insert) and smell (through use of a mild essence dilution added to the cleaning solution). Video scoring is automated, and occurs in real-time, by the proprietary Video Freeze (Med Associates Inc) software package, with freezing behavior programmed as a lack of movement detected across 30 frames (one second). On Day 1, animals underwent a 10 min trial period with a 3.5 min baseline period followed by 5-foot shocks of 0.9 mA intensity paired with an auditory tone (90 dB, 5 kHz). The auditory tone (5 s duration) preceded the foot shocks (200 ms duration). Equipment was cleansed with 70% ethanol between animal trials. On Day 2, mice were placed in the same context chamber as Day 1 and freezing behavior was calculated for a trial period of 5 min. Equipment was cleansed with 70% ethanol across trials. On Day 3, white plexi-glass inserts were placed in the test chamber to simulate a novel context environment. During the 5 min trial period 3 auditory tones were delivered to test cued memory.

### Chemicals

Chemicals for ACSF, NMDG, sucrose (S0389), and picrotoxin (P1675) were obtained from Sigma-Aldrich. NMDG (M2004), KCl (P3911), NaH_2_PO_4_ (S9638), NaHCO_3_ (S6014), HEPES (H3375), glucose (G8270), sodium ascorbate (A4034), Thiourea (T8656), sodium pyruvate (P2256), MgCl_2_ (M2670), CaCl_2_ (C8106), NaCl (S7653), MgSO_4_ (M1880). DHPG (0342) and CGP-55845 (1248), was ordered from Tocris Bioscience. HCl (SA49) was obtained from Fisher Scientific.

### Statistical Analysis

Statistical Analyses and graphical data formatting were performed using GraphPad Prism 10 or Origin Pro 8 software. All data distributions were tested for normality using the Shapiro-Wilk test. Normal data distributions underwent parametric Unpaired *t*-test analysis for sample sizes (N > 7). For skewed distributions or sample sizes (N < 7) the non-parametric Mann-Whitney test was applied. For behavioral experiments, the Sidak’s multiple comparison test was implemented for Day 1 of the contextual fear conditioning paradigm. The CalEx rescue experiments were analyzed using One-way ANOVA followed by post-hoc Tukey’s multiple comparisons test when statistical significant interaction was detected. Statistical significance was determined for p values < 0.05 associated with the statistical analysis test applied. All experimental data collection and analysis was performed in a blind-manner.

## Supporting information

CTRL Astrocyte spontaneous calcium events

Upf2 cKO Astrocyte spontaneous calcium events

CalEx restoration of calcium events in cKO astrocytes

## RESOURCE AVAILABILITY

### Lead contact

Further information and requests for resources should be directed to the lead contact, Dr. Dilek Colak.

### Materials availability

All materials are available commercially, as detailed in the Materials and Methods. All data associated with this study are in the main text or the supplemental information.

### Data and code availability

RNA sequencing data will be uploaded to dbGAP upon publication. This study does not report original code. Information required to reanalyze data reported is available from the lead contact upon request.

## ACKNOWLEDGEMENTS

We thank the Weill Cornell Medicine Genomics and Fluorescence Activated Cell Sorting Core facilities for providing experimental consultation, assistance and results. We also thank Jens Lykke-Andersen for generously providing the anti-UPF2 antibody. We thank Natalie Wayland and Kelsey Lysek for support maintaining the mouse colonies and Mars Reynoso for performing pilot immunostainings.

## AUTHOR CONTRIBUTIONS

D.C. and P.J.L conceived of the project, designed experiments, and wrote the manuscript with input from all authors. P.J.L performed majority of the experiments including: astrocyte cell selection, electrophysiology, calcium imaging, stereotaxic surgeries, behavioral tests, immunostainings, Imaris analyses, and executed associated analyses. A.D. performed Imaris analyses and associated immunostainings. E.B.A. supported behavioral tests. K.T. performed RNA-seq and NMD inducing feature analyses with supervision from M.W. Support for two-photon microscopy experiments was provided by P.E.C.

## DECLARATION OF GENERATIVE AI

Generative AI was not implemented in the writing or analytical aspects of the study.

