## Supplementary Figures for "Astrocytic Nonsense-mediated mRNA decay regulates calcium signaling to support synapse function and influence anxiety-like behavior"

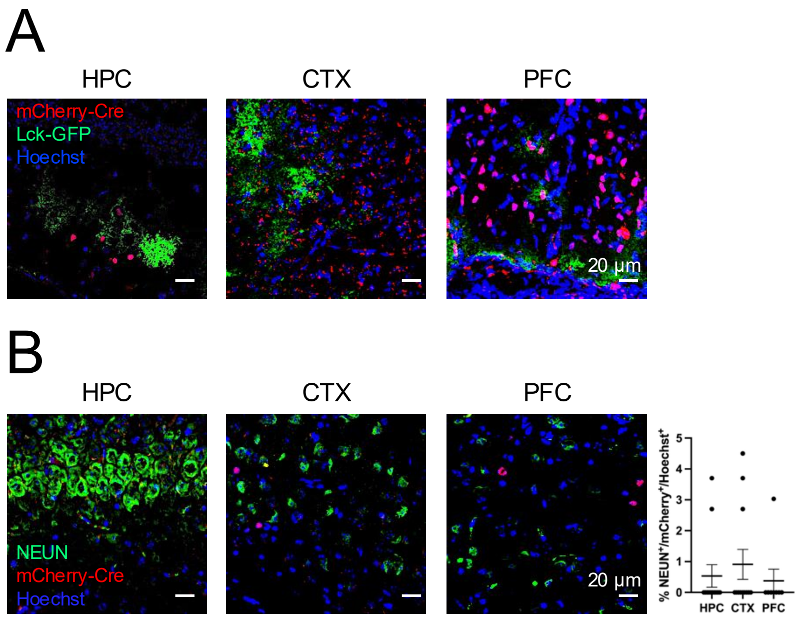


**Figure S1. Viral expression of mCherry-Cre and Lck-GFP reporters across brain areas.** (**A**) Images depicting region of interest (ROI) views of mCherry-Cre$^{+}$ and Lck-GFP$^{+}$ expression in hippocampus, visual cortex, and prefrontal cortex. (**B**) Images showing negligible NEUN/mCherry colocalization across targeted brain areas (HPC: 0.5 ± 0.4 %, n = 6 animals, 2 FOVs per animal, CTX: 0.9 ± 0.5 %, 6 animals, 2 FOVs per animal, PFC: 0.4 ± 0.4 %, n = 4 animals, 2 FOVs per animal). Data is presented as mean ± S.E.M. **This figure highlights that mCherry-Cre and Lck-GFP viruses are expressed in targeted brain areas with negligible neuronal leak.**

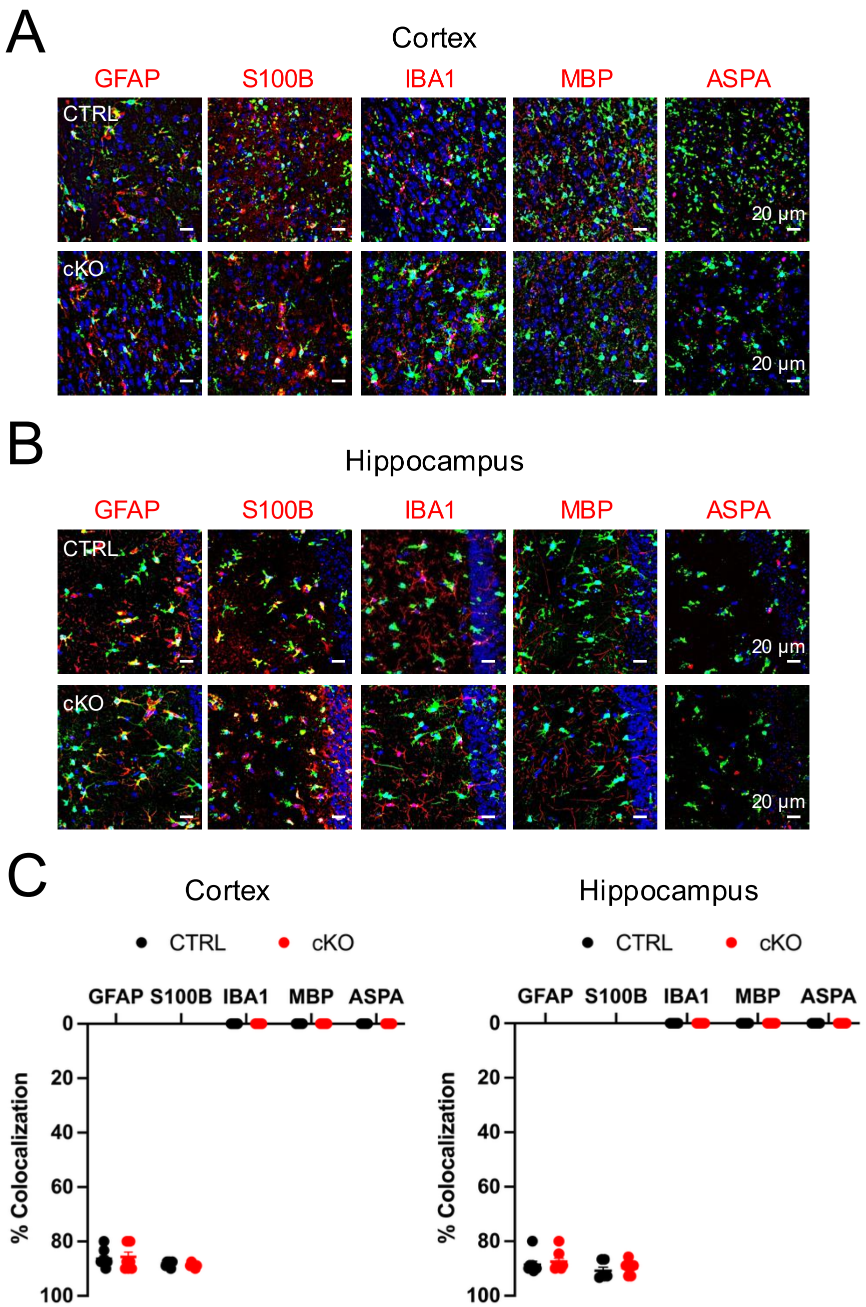


**Figure S2. ZsGreen1 selectively colocalizes with canonical astrocyte markers GFAP and S100B.** In all panels of this figure canonical makers for astrocytes (GFAP, S100B), microglia (IBA1) and oligodendrocytes (MBP, ASPA) are shown in red. (**A-B**) Cortical and hippocampal images depicting colocalization of GFAP$^{+}$ and S100B$^{+}$ astrocytes with ZsGreen1 as compared to IBA1, MBP, and ASPA in CTRL and cKO animals. (**C**) Quantification reveals high colocalization of ZsGreen1 with canonical astrocyte markers in cortex of both genotypes (GFAP; CTRL: 86 ± 1%, cKO: 86 ± 2%; S100B; CTRL: 88 ± 0.4%, cKO: 88 ± 0.3%, n = 3-4 mice, 2 FOVs per animal). Hippocampal quantification also demonstrates high colocalization of ZsGreen1 with astrocyte markers across groups (GFAP; CTRL: 88 ± 1%, cKO: 87 ± 1%; S100B; CTRL: 91 ± 1%, cKO: 90 ± 1%, n = 3-4 mice, 2 FOVs per animal). Data is presented as mean ± S.E.M. **This figure indicates the selective expression of ZsGreen1 reporter protein in astrocytes of the cortex and hippocampus.**

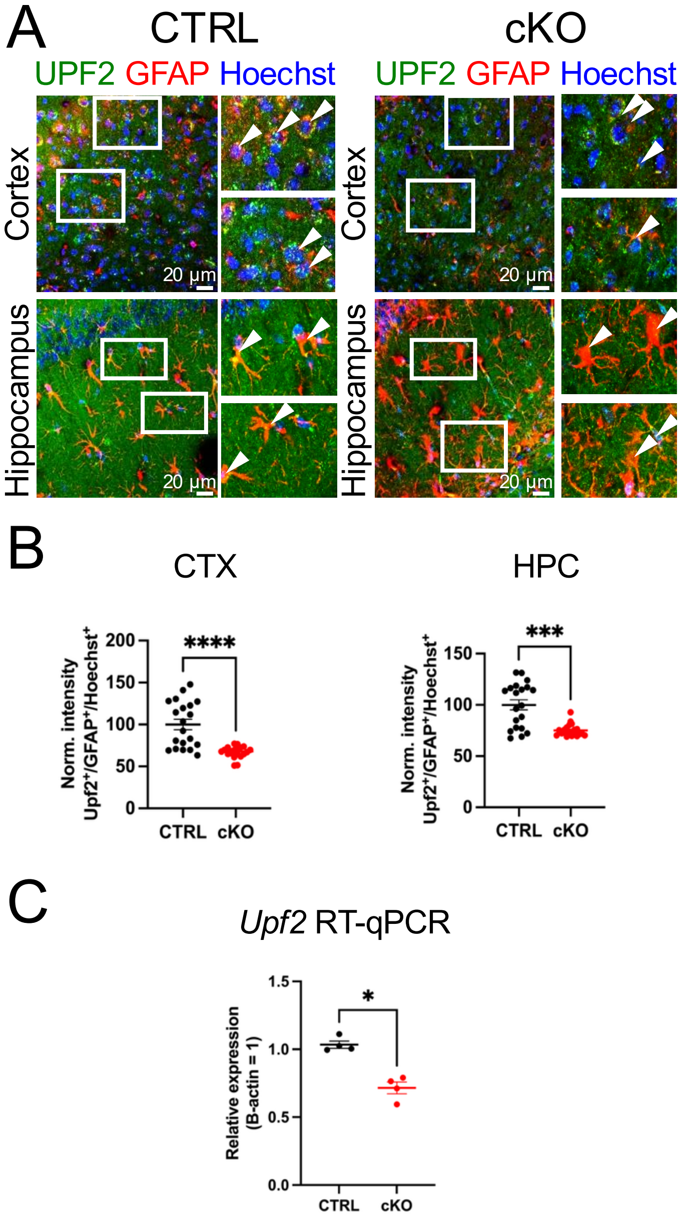


**Figure S3. UPF2 protein and transcript levels are reduced in cKO conditions.** (**A**) Confocal microscopy images depicting regions of interest (ROI, white boxes) highlighting UPF2$^+$/GFAP$^+$ astrocytes in hippocampus and cortex. (**B**) UPF2 protein is reduced in cKO mice (CTX: CTRL: 100 ± 6, cKO: 67 ± 2, p < 0.0001, Unpaired *t*-test; HPC: CTRL: 100 ± 5, cKO: 75 ± 1, p = 0.0003, Mann-Whitney test, n = 4 mice per genotype, 20 ROIs per group). (**C**) RT-qPCR of ZsGreen1$^+$ CTRL and cKO astrocytes reveals a significant reduction of *Upf2* mRNA (CTRL: 1.03 ± 0.03, cKO: 0.72 ± 0.04, n = 4 cell selections from 3-4 mice per genotype, p = 0.0286, Mann-Whitney test). Data is presented as mean ± S.E.M. *** denotes p < 0.005, **** denotes p < 0.001 * denotes p < 0.05. **This figure confirms the downregulation of UPF2 protein and *Upf2* transcript levels in cKO mice.**

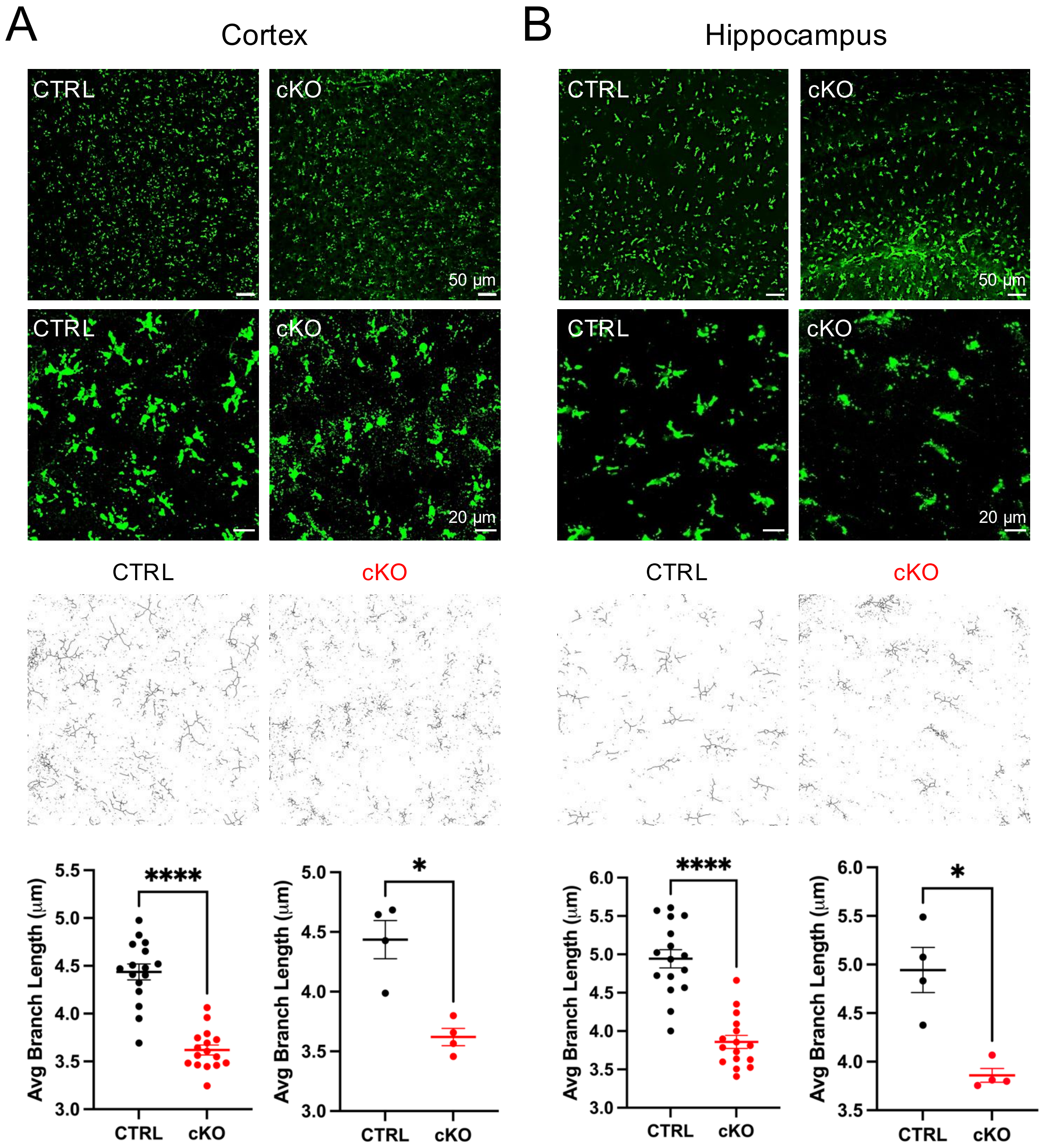


**Figure S4. *Upf2* loss alters astrocyte cell size by reducing branch length.** (**A**) Representative low and high magnification images of cortical astrocytes in CTRL and cKO mice. Average branch length quantification using skeletonization demonstrates a significant reduction in *Upf2* cKO astrocyte cell size as compared to control (CTRL: 4.4 ± 0.08 cKO: 3.6 ± 0.05, n = 4 animals, 4 FOVs per animal, $p < 0.0001$, Unpaired *t*-test). (**B**) Representative low and high magnification images of hippocampal astrocytes in CTRL and cKO mice. Average branch length using skeletonization reveals a significant reduction in *Upf2* cKO astrocyte cell size as compared to control (CTRL: 4.9 ± 0.12 cKO: 3.8 ± 0.09, n = 4 animals, 4 FOVs per animal, $p < 0.0001$, Unpaired *t*-test). Summary data for each brain region illustrating average branch length per animal (CTRL: 4.4 ± 0.16 CTX, 4.9 ± 0.2 HPC; cKO: 3.6 ± 0.07, CTX, 3.8 ± 0.07, HPC, n = 4 animals per genotype group, p = 0.028, Mann-Whitney test). Data is presented as mean ± S.E.M. **** denotes $p < 0.0001$, * denotes $p < 0.05$. **This figure indicates astrocyte branch length is reduced contributing to smaller cell size in *Upf2* cKO conditions.**

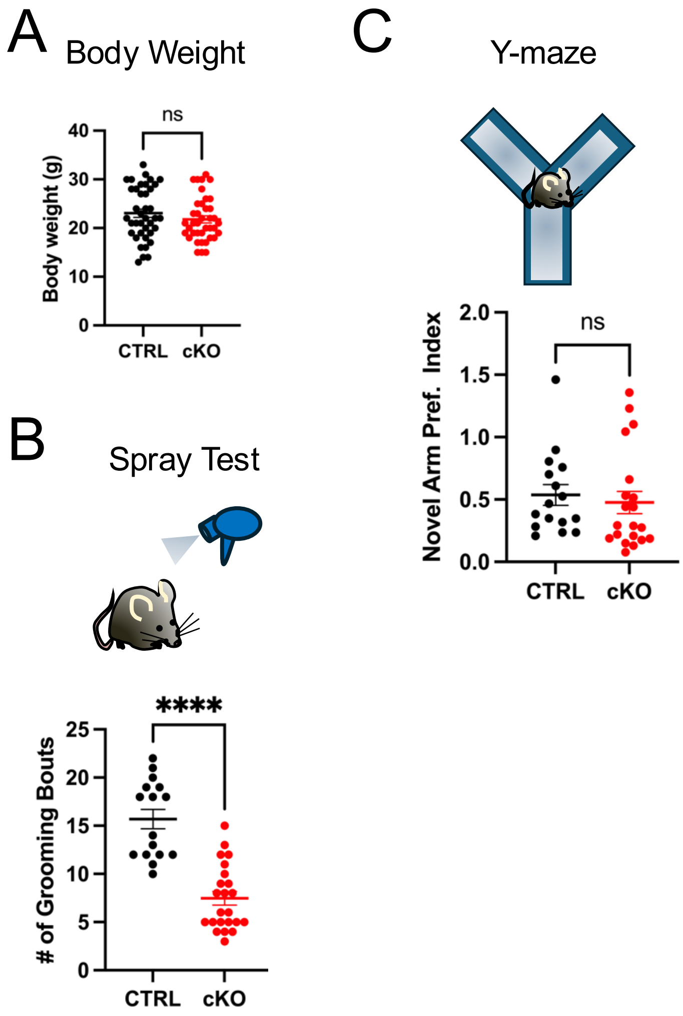


**Figure S5. *Upf2* cKO mice display normal body weight, reduced grooming, and intact short-term memory.** (**A**) *Upf2* cKO and CTRL animals exhibit similar body weight (CTRL: 23.1 ± 0.9, cKO: 21.8 ± 0.7, n = 39 mice per group, p = 0.24, Mann-Whitney test). (**B**) *Upf2* cKO mice groomed less in response to water spray as compared to control (CTRL: 15.7 ± 1, n = 16 mice; CKO: 7.5 ± 0.7, n = 23 mice, p < 0.0001, Mann-Whitney test). (**C**) CTRL and cKO animals show similar novel arm preference in the Y-maze indicating intact short-term memory (CTRL: 0.5 ± 0.08, n = 16, cKO: 0.48 ± 0.09, n = 20 mice, p = 0.18, Mann-Whitney test). Data are presented as mean ± S.E.M. **** denotes p < 0.0001, n.s. = not significant. **This figure illustrates *Upf2* cKO mice groom less and display intact short-term memory.**

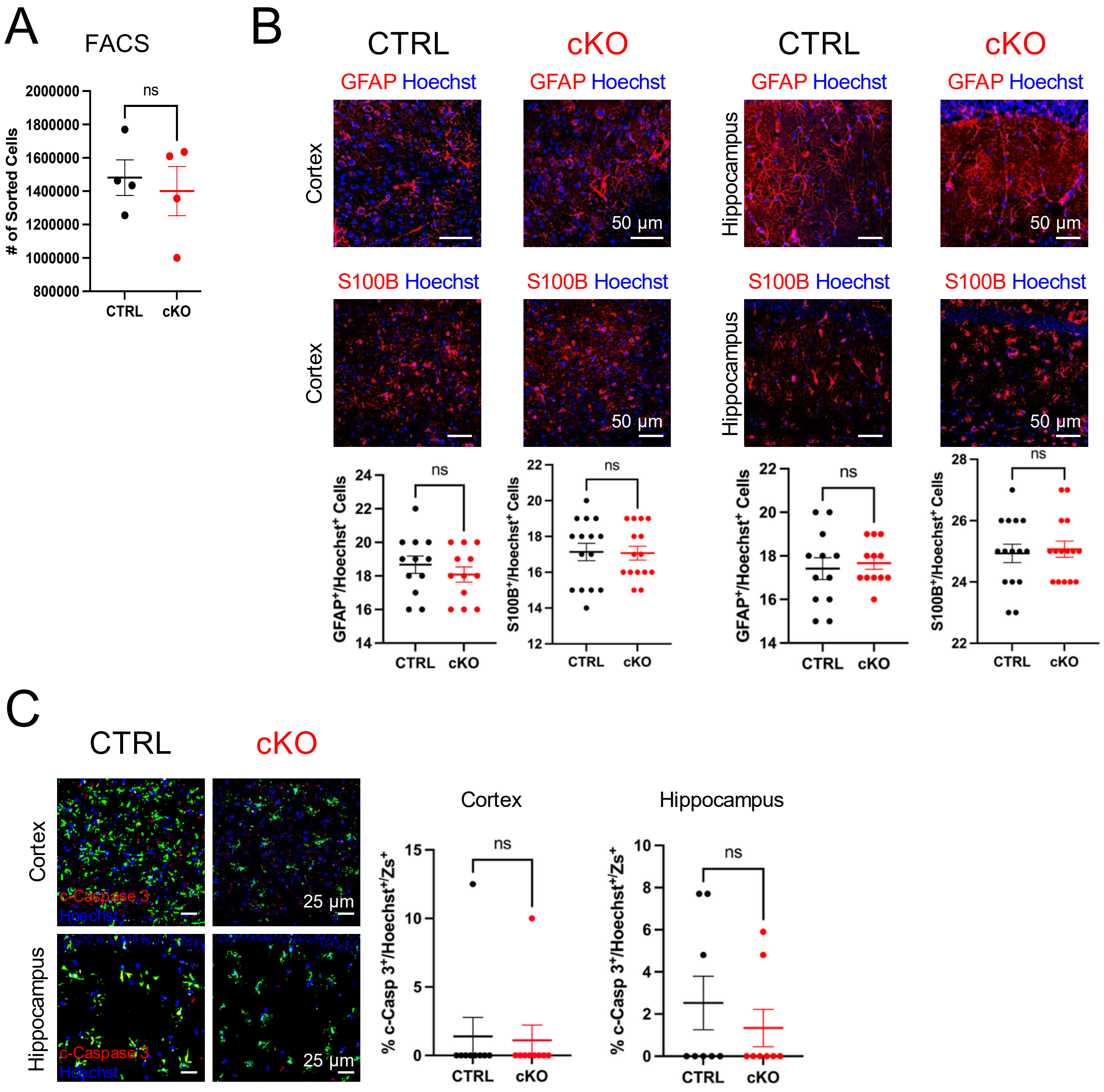


**Figure S6. *Upf2* loss does not change astrocyte cell number or cell survival.** (**A**) FACS procedures revealed no differences in cell number (CTRL: 1.48 x $10^6$ ± 0.11, cKO: 1.40 x $10^6$ ± 0.15, 4 technical replicates, 3 mice per genotype per replicate, p = 0.88, Mann-Whitney test). (**B**) No difference in astrocyte cell number is shown across genotypes in cortex and hippocampus immunostaining with GFAP and S100B, respectively (GFAP CTX: CTRL: 18.6 ± 0.5, cKO: 18.1 ± 0.5, p = 0.47, Mann-Whiteny test; GFAP HPC: CTRL: 17.4 ± 0.5, cKO: 17.6 ± 0.3, p = 0.69, Mann-Whiteny test; n = 4 mice, 3 FOVs per animal; S100B CTX: CTRL: 17.1 ± 0.5, cKO: 17 ± 0.4, p = 0.96, Mann-Whiteny test; S100B HPC: CTRL: 24.9 ± 0.3, cKO: 25 ± 0.3, p = 0.91, Mann-Whiteny test; n = 5 mice, 3 FOVs per animal). (**C**) Quantification of nuclear cleaved-Caspase 3 reveals no differences in cortex and hippocampus indicating comparable apoptosis in CTRL and cKO animals (CTX: CTRL: 1.4 ± 1.4, cKO: 1.1 ± 1.1, p > 0.9, Mann-Whiteny test; HPC: CTRL: 2.5 ± 1.2, cKO: 1.3 ± 0.8, p = 0.49, Mann-Whiteny test; n = 3 mice, 3 FOVs per animal). Data is presented as mean ± S.E.M. ns = not significant. **This figure indicates astrocyte cell number and cell survival is unchanged in cortex and hippocampus of *Upf2* cKO animals.**

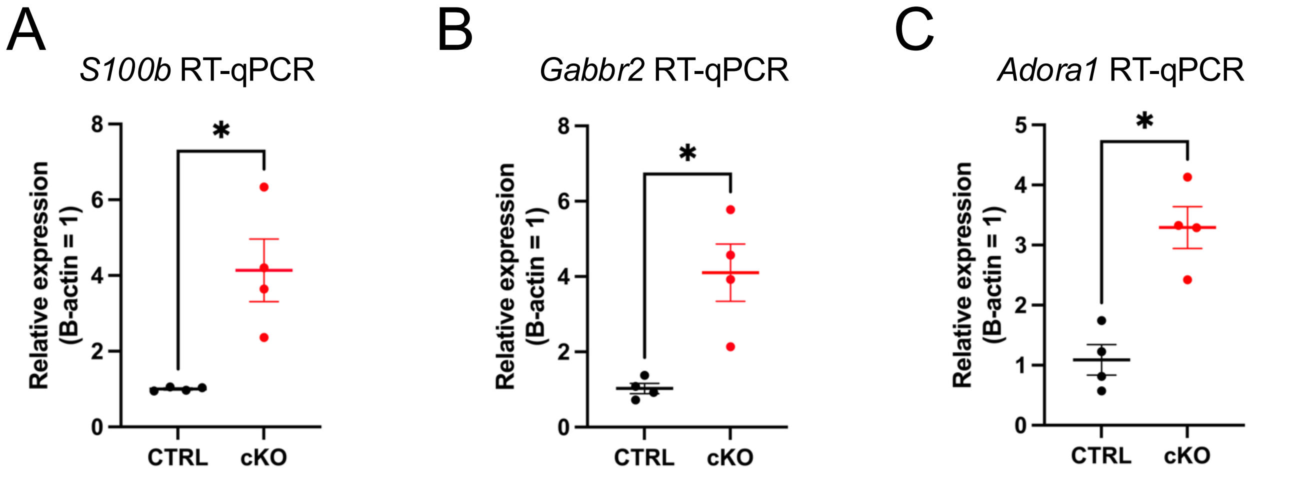


**Figure S7. Validation of RNA-seq astrocyte NMD targets with RT-qPCR.** Total RNA extracted from CTRL and cKO FACS sorted ZsGreen1$^{+}$ astrocytes enabled cDNA synthesis for RT-qPCR using the SYBR method. Comparative $C_t$ revealed a significant upregulation of candidate NMD targets, *S100b*, *Gabbr2*, *and Adora1.* In this experiment, target genes were normalized to housekeeping gene *B-actin.* (**A**) RT-qPCR of ZsGreen1$^{+}$ astrocytes shows a significant *S100b* upregulation in cKO astrocytes as compared to control (CTRL: 1.0 ± 0.02, cKO: 4.1 ± 0.8, n = 4 cell selections from 3-4 mice per genotype, p = 0.0286, Mann-Whitney test). (**B**) RT-qPCR of ZsGreen1$^{+}$ astrocytes indicated *Gabbr2* upregulation in cKO conditions as compared to control (CTRL: 1.03 ± 0.14, cKO: 4.1 ± 0.8, n = 4 cell selections from 3-4 mice per genotype, p = 0.0286, Mann-Whitney test). (**C**) RT-qPCR of ZsGreen1$^{+}$ astrocytes demonstrate a significant *Adora1* upregulation in cKO conditions in comparison to control (CTRL: 1.09 ± 0.26, cKO: 3.3 ± 0.35, n = 4 cell selections from 3-4 mice per genotype, p = 0.0286, Mann-Whitney test). Data is presented as mean ± S.E.M. * denotes $p < 0.05$. **This figure supports the finding that *S100b*, *Gabbr2*, and *Adora1* are upregulated astrocyte transcripts in *Upf2* cKO conditions.**

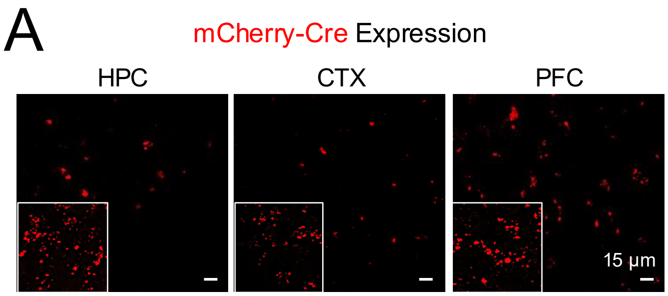


**Figure S8. Viral expression of mCherry-Cre reporter across brain areas.** (**A**) Two-photon images depicting region of interest (ROI) views of mCherry-Cre reporter expression in hippocampus, visual cortex, and prefrontal cortex in acute coronal brain slices. Box inserts show zoom-in of ROIs. **This figure highlights that mCherry-Cre virus is expressed in targeted brain areas.**

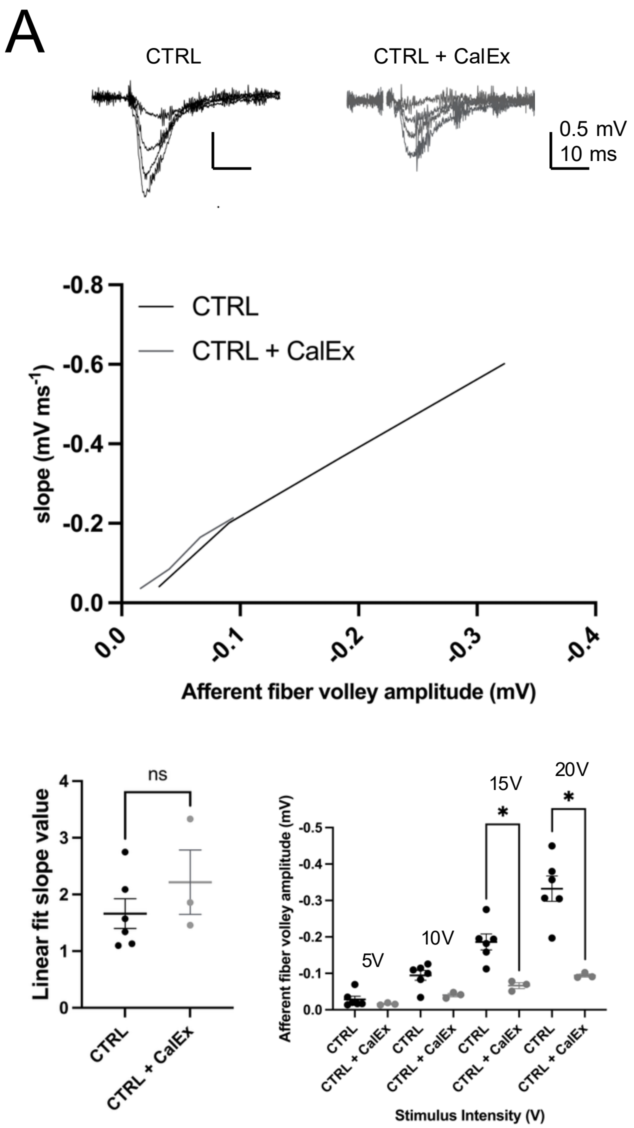


**Figure S9. CalEx expression in CTRL animals does not impact linear-fit slope values but minimizes fiber volley amplitude.** (**A**) Representative traces of fEPSP responses from CTRL and CTRL + CalEx acute hippocampal slices *(top)*. Summary plot indicating a shift in fiber volley amplitude and fEPSP slope (mV $ms^{-1}$) in CTRL + CalEx hippocampal slices as compared to CTRL with no major changes to linear regression values, CTRL: Linear fit 1.88 ± 0.1, r = 0.99; CTRL + CalEx: Linear fit 2.2 ± 0.2, r = 0.98 *(middle)*. Summary data of linear fit slope values revealing no significant differences in CTRL and CTRL + CalEx mice (*bottom*, *left:* CTRL: 1.66 ± 0.3, n = 5 slices, 3 mice; CTRL + CalEx: 2.2 ± 0.6, n = 3 slices, 3 mice, p = 0.38, Mann-Whitney test). CalEx significantly minimizes fiber volley amplitude in CTRL animals (*bottom*, *right*: 15V, CTRL: -0.19 ± 0.02, n = 5 slices, 3 mice; CTRL + CalEx: -0.07 ± 0.008, n = 3 slices, 3 mice, p = 0.0238, Mann-Whitney test; 20V, CTRL: -0.33 ± 0.03, n = 5 slices, 3 mice; CTRL + CalEx: -0.09 ± 0.004, n = 3 slices, 3 mice, p < 0.0238, Mann-Whitney test). Data is presented as mean ± S.E.M. * denotes p < 0.005, ns = not significant. **This figure shows that while CalEx does not significantly alter linear fit slope values, CalEx decreases fiber volley amplitude in CTRL animals.**
