## Supplemental Table S1 for "Astrocytic Nonsense-mediated mRNA decay regulates calcium signaling to support synapse function and influence anxiety-like behavior"

**Table S1. Upregulated and Downregulated DEG List**

| gene_id | baseMean | log2FoldCh | lfcSE | stat | pvalue | padj | Gene_symbol |
|---|---|---|---|---|---|---|---|
| ENSMUSG00000086231 | 54.40342 | 9.234591 | 1.574111 | 5.866543 | 4.45E-09 | 4.56E-06 | NA |
| ENSMUSG00000036853 | 21.1604 | 7.872299 | 1.515823 | 5.193415 | 2.06E-07 | 8.19E-05 | Mcoln3 |
| ENSMUSG00000026390 | 393.486 | 5.824038 | 1.521349 | 3.828206 | 0.000129 | 0.011625 | Marco |
| ENSMUSG00000079304 | 54.44421 | 5.216555 | 1.006649 | 5.182099 | 2.19E-07 | 8.53E-05 | Tex52 |
| ENSMUSG00000033383 | 53.01663 | 5.131662 | 0.726407 | 7.064445 | 1.61E-12 | 8.00E-09 | Rtp1 |
| ENSMUSG00000032517 | 87242.46 | 4.872026 | 0.973346 | 5.00544 | 5.57E-07 | 0.000177 | Mobp |
| ENSMUSG00000094420 | 47.05706 | 4.863244 | 1.422614 | 3.418527 | 0.00063 | 0.035216 | Igkv10-96 |
| ENSMUSG00000000791 | 48.25457 | 4.824388 | 0.740015 | 6.519314 | 7.06E-11 | 1.56E-07 | Il12rb1 |
| ENSMUSG00000040026 | 180.7948 | 4.676933 | 0.7417 | 6.305692 | 2.87E-10 | 5.17E-07 | Saa3 |
| ENSMUSG00000041607 | 296615.3 | 4.614964 | 0.719599 | 6.413245 | 1.42E-10 | 2.83E-07 | Mbp |
| ENSMUSG00000006777 | 89.87295 | 4.529404 | 0.930277 | 4.868878 | 1.12E-06 | 0.000299 | Krt23 |
| ENSMUSG00000013523 | 6534.758 | 4.44778 | 0.972596 | 4.573102 | 4.81E-06 | 0.000983 | Bcas1 |
| ENSMUSG00000105843 | 179.1286 | 4.419033 | 0.653931 | 6.757641 | 1.40E-11 | 3.97E-08 | NA |
| ENSMUSG00000023943 | 101.641 | 4.415158 | 0.991086 | 4.454869 | 8.39E-06 | 0.001571 | Sult1c1 |
| ENSMUSG00000044177 | 63.18239 | 4.270315 | 0.99964 | 4.271855 | 1.94E-05 | 0.002785 | Wfikkn2 |
| ENSMUSG00000104093 | 84.12528 | 4.227218 | 1.189613 | 3.553439 | 0.00038 | 0.025615 | A330015K06Rik |
| ENSMUSG00000036687 | 45.6997 | 4.214528 | 0.966312 | 4.361459 | 1.29E-05 | 0.002118 | Tmem184a |
| ENSMUSG00000074796 | 41.69672 | 4.141198 | 0.828989 | 4.995482 | 5.87E-07 | 0.000179 | Slc4a11 |
| ENSMUSG00000047109 | 18.23497 | 4.096179 | 1.085797 | 3.772509 | 0.000162 | 0.013584 | Cldn14 |
| ENSMUSG00000055945 | 3364.962 | 4.08667 | 0.668533 | 6.112891 | 9.78E-10 | 1.29E-06 | Prr18 |
| ENSMUSG00000041460 | 24.0443 | 3.992403 | 0.918297 | 4.347617 | 1.38E-05 | 0.002204 | Cacna2d4 |
| ENSMUSG00000017734 | 1506.905 | 3.946589 | 0.651387 | 6.058746 | 1.37E-09 | 1.70E-06 | Dbndd2 |
| ENSMUSG00000104494 | 36.28944 | 3.870252 | 0.985553 | 3.926985 | 8.60E-05 | 0.008618 | NA |
| ENSMUSG00000002588 | 51.22169 | 3.815763 | 0.947537 | 4.027035 | 5.65E-05 | 0.006157 | Pon1 |
| ENSMUSG00000023122 | 188.3144 | 3.799514 | 0.620571 | 6.122611 | 9.21E-10 | 1.29E-06 | Sult1c2 |
| ENSMUSG00000040170 | 160.3661 | 3.729683 | 0.962607 | 3.874562 | 0.000107 | 0.010286 | Fmo2 |
| ENSMUSG00000015401 | 83.46812 | 3.661206 | 0.962238 | 3.804887 | 0.000142 | 0.012338 | Cltrn |
| ENSMUSG00000030701 | 22955.27 | 3.606334 | 0.670835 | 5.375886 | 7.62E-08 | 4.64E-05 | Plekhb1 |
| ENSMUSG00000078137 | 22.70324 | 3.546204 | 0.895907 | 3.958229 | 7.55E-05 | 0.007721 | Ankrd63 |
| ENSMUSG00000104057 | 19.35595 | 3.468862 | 0.78437 | 4.42248 | 9.76E-06 | 0.001776 | NA |
| ENSMUSG00000053846 | 31.59142 | 3.464648 | 0.957777 | 3.617384 | 0.000298 | 0.021235 | Lipg |
| ENSMUSG00000107451 | 120.929 | 3.400508 | 0.649534 | 5.235305 | 1.65E-07 | 7.23E-05 | NA |
| ENSMUSG00000025500 | 20.22858 | 3.395771 | 1.012864 | 3.352642 | 0.0008 | 0.041566 | Lmntd2 |
| ENSMUSG00000110027 | 70.07685 | 3.371119 | 0.783179 | 4.304403 | 1.67E-05 | 0.002555 | C030029H02Rik |
| ENSMUSG00000030638 | 1019.806 | 3.363538 | 0.676939 | 4.968746 | 6.74E-07 | 0.000194 | Sh3gl3 |
| ENSMUSG00000039601 | 3317.431 | 3.223236 | 0.607074 | 5.309459 | 1.10E-07 | 5.59E-05 | Rcan2 |
| ENSMUSG00000004894 | 268.787 | 3.21771 | 0.443472 | 7.255716 | 4.00E-13 | 2.64E-09 | Hapln2 |
| ENSMUSG00000071561 | 54.42511 | 3.194511 | 0.583555 | 5.474227 | 4.39E-08 | 3.10E-05 | Cstdc5 |
| ENSMUSG00000020427 | 769.5876 | 3.180567 | 0.639232 | 4.975605 | 6.50E-07 | 0.000193 | Igfbp3 |
| ENSMUSG00000071562 | 63.64571 | 3.093434 | 0.453474 | 6.821643 | 9.00E-12 | 2.98E-08 | Stfa1 |
| ENSMUSG00000079594 | 29.97934 | 3.055672 | 0.716208 | 4.266461 | 1.99E-05 | 0.002794 | Cstdc6 |
| ENSMUSG00000100147 | 66.99924 | 3.03935 | 0.463459 | 6.557966 | 5.45E-11 | 1.35E-07 | 1700047M11Rik |
| ENSMUSG00000098682 | 62.47508 | 3.007782 | 0.686557 | 4.380965 | 1.18E-05 | 0.002003 | Otx2os1 |
| ENSMUSG00000104674 | 54.93009 | 2.914503 | 0.663691 | 4.391357 | 1.13E-05 | 0.00196 | NA |
| ENSMUSG00000041309 | 588.9861 | 2.908844 | 0.51007 | 5.702833 | 1.18E-08 | 1.02E-05 | Nkx6-2 |
| ENSMUSG00000022902 | 35.58436 | 2.88845 | 0.539095 | 5.357957 | 8.42E-08 | 4.64E-05 | Stfa2 |

| | | | | | | | |
|---|---|---|---|---|---|---|---|
| ENSMUSG00000030878 | 565.984 | 2.178747 | 0.59165 | 3.682493 | 0.000231 | 0.01769 | Cdr2 |
| ENSMUSG00000027858 | 3550.14 | 2.174514 | 0.471852 | 4.608469 | 4.06E-06 | 0.000838 | Tspan2 |
| ENSMUSG00000039661 | 1128.23 | 2.172352 | 0.492926 | 4.407054 | 1.05E-05 | 0.00187 | Dusp26 |
| ENSMUSG00000037625 | 355.6988 | 2.164347 | 0.504331 | 4.291519 | 1.77E-05 | 0.002627 | Cldn11 |
| ENSMUSG00000111013 | 57.34951 | 2.160853 | 0.512845 | 4.21346 | 2.51E-05 | 0.003364 | NA |
| ENSMUSG00000060716 | 824.3026 | 2.148006 | 0.493966 | 4.348491 | 1.37E-05 | 0.002204 | Plekhh1 |
| ENSMUSG00000056162 | 54.16044 | 2.135027 | 0.543204 | 3.930436 | 8.48E-05 | 0.008582 | Cndp1 |
| ENSMUSG00000028111 | 73.54639 | 2.121313 | 0.591263 | 3.587768 | 0.000334 | 0.023214 | Ctsk |
| ENSMUSG00000038007 | 1101.595 | 2.111489 | 0.549861 | 3.840044 | 0.000123 | 0.011245 | Acer2 |
| ENSMUSG00000061808 | 129526.1 | 2.10464 | 0.509419 | 4.131454 | 3.60E-05 | 0.004387 | Ttr |
| ENSMUSG00000056222 | 267.4925 | 2.102446 | 0.461685 | 4.553858 | 5.27E-06 | 0.001055 | Spock1 |
| ENSMUSG00000061451 | 1015.402 | 2.075448 | 0.438208 | 4.736217 | 2.18E-06 | 0.000505 | Tmem151a |
| ENSMUSG00000076439 | 1743.361 | 2.074251 | 0.447996 | 4.630069 | 3.66E-06 | 0.00078 | Mog |
| ENSMUSG00000004655 | 962.7369 | 2.064625 | 0.53219 | 3.879489 | 0.000105 | 0.010229 | Aqp1 |
| ENSMUSG00000026904 | 1100.097 | 2.051235 | 0.535937 | 3.827379 | 0.00013 | 0.011625 | Slc4a10 |
| ENSMUSG00000010080 | 597.8242 | 2.026764 | 0.391486 | 5.177101 | 2.25E-07 | 8.60E-05 | Epn3 |
| ENSMUSG00000049721 | 349.9163 | 2.015362 | 0.598037 | 3.369962 | 0.000752 | 0.039938 | Gal3st1 |
| ENSMUSG00000097365 | 31.73024 | 1.995882 | 0.601472 | 3.318329 | 0.000906 | 0.045826 | C030034L19Rik |
| ENSMUSG00000064325 | 165.7794 | 1.99453 | 0.452744 | 4.405426 | 1.06E-05 | 0.00187 | Hhip |
| ENSMUSG00000030683 | 794.286 | 1.993918 | 0.44921 | 4.438723 | 9.05E-06 | 0.001662 | Sez6l2 |
| ENSMUSG00000073600 | 211.2144 | 1.989942 | 0.556246 | 3.577449 | 0.000347 | 0.024065 | Prob1 |
| ENSMUSG00000117313 | 213.2051 | 1.988068 | 0.478861 | 4.15166 | 3.30E-05 | 0.004092 | NA |
| ENSMUSG00000030450 | 208.4631 | 1.98232 | 0.511499 | 3.875513 | 0.000106 | 0.010286 | Oca2 |
| ENSMUSG00000032186 | 3257.324 | 1.974921 | 0.516606 | 3.82288 | 0.000132 | 0.011681 | Tmod2 |
| ENSMUSG00000047502 | 60.13844 | 1.973084 | 0.532893 | 3.702591 | 0.000213 | 0.016472 | Mroh7 |
| ENSMUSG00000084289 | 3464.697 | 1.965065 | 0.480706 | 4.087871 | 4.35E-05 | 0.00508 | NA |
| ENSMUSG00000086284 | 67.68858 | 1.960584 | 0.459114 | 4.270365 | 1.95E-05 | 0.002785 | Frmpd1os |
| ENSMUSG00000028125 | 816.2211 | 1.957839 | 0.521809 | 3.752019 | 0.000175 | 0.014499 | Abca4 |
| ENSMUSG00000001348 | 419.9368 | 1.945754 | 0.37625 | 5.171442 | 2.32E-07 | 8.69E-05 | Acp5 |
| ENSMUSG00000015090 | 13234.58 | 1.935535 | 0.438681 | 4.412171 | 1.02E-05 | 0.001846 | Ptgds |
| ENSMUSG00000020812 | 164.1893 | 1.93161 | 0.405351 | 4.765279 | 1.89E-06 | 0.000462 | Snhg16 |
| ENSMUSG00000024661 | 141019.1 | 1.919618 | 0.459174 | 4.180587 | 2.91E-05 | 0.003721 | Fth1 |
| ENSMUSG00000025584 | 625.2351 | 1.91947 | 0.406867 | 4.717688 | 2.39E-06 | 0.000538 | Pde8a |
| ENSMUSG00000029843 | 1020.237 | 1.91785 | 0.552703 | 3.469951 | 0.000521 | 0.031871 | Slc13a4 |
| ENSMUSG00000091735 | 248.3456 | 1.909223 | 0.530964 | 3.595768 | 0.000323 | 0.022671 | Gpr62 |
| ENSMUSG00000066720 | 92.20441 | 1.893985 | 0.539538 | 3.510386 | 0.000447 | 0.028412 | Cldn9 |
| ENSMUSG00000024810 | 1652.856 | 1.888353 | 0.318106 | 5.936242 | 2.92E-09 | 3.21E-06 | Il33 |
| ENSMUSG00000033579 | 881.4329 | 1.886221 | 0.431776 | 4.368514 | 1.25E-05 | 0.002103 | Fa2h |
| ENSMUSG00000028789 | 699.2824 | 1.883805 | 0.36452 | 5.167903 | 2.37E-07 | 8.70E-05 | Azin2 |
| ENSMUSG00000042349 | 390.8505 | 1.879026 | 0.455655 | 4.123792 | 3.73E-05 | 0.004508 | Ikbke |
| ENSMUSG00000032373 | 1393.061 | 1.873386 | 0.457027 | 4.099073 | 4.15E-05 | 0.004927 | Car12 |
| ENSMUSG00000020774 | 621.6149 | 1.873098 | 0.52827 | 3.54572 | 0.000392 | 0.026064 | Aspa |
| ENSMUSG00000024867 | 246.7162 | 1.873088 | 0.363125 | 5.158242 | 2.49E-07 | 8.99E-05 | Pip5k1b |
| ENSMUSG00000015653 | 619.1141 | 1.866521 | 0.508212 | 3.672719 | 0.00024 | 0.018059 | Steap2 |
| ENSMUSG00000038375 | 6105.414 | 1.866167 | 0.48083 | 3.881133 | 0.000104 | 0.01021 | Trp53inp2 |
| ENSMUSG00000033595 | 1888.592 | 1.86442 | 0.416511 | 4.476275 | 7.60E-06 | 0.001449 | Lgi3 |
| ENSMUSG00000045087 | 1172.229 | 1.861511 | 0.32212 | 5.77893 | 7.52E-09 | 7.10E-06 | S1pr5 |

| | | | | | | | |
|---|---|---|---|---|---|---|---|
| ENSMUSG00000038668 | 2215.228 | 2.885318 | 0.664488 | 4.342168 | 1.41E-05 | 0.002239 | Lpar1 |
| ENSMUSG00000117239 | 102.1308 | 2.873214 | 0.654 | 4.39329 | 1.12E-05 | 0.00196 | Gpr31c |
| ENSMUSG00000026418 | 134.9585 | 2.809872 | 0.652149 | 4.308637 | 1.64E-05 | 0.002531 | Tnni1 |
| ENSMUSG00000002831 | 137.0719 | 2.797483 | 0.786869 | 3.555208 | 0.000378 | 0.02557 | Plin4 |
| ENSMUSG00000117864 | 53.62903 | 2.795296 | 0.789968 | 3.538494 | 0.000402 | 0.026609 | Gm19500 |
| ENSMUSG00000101304 | 49.09432 | 2.790055 | 0.67964 | 4.105197 | 4.04E-05 | 0.004827 | Plet1os |
| ENSMUSG00000028584 | 48.98229 | 2.779206 | 0.809793 | 3.431995 | 0.000599 | 0.034586 | Lrrc38 |
| ENSMUSG00000116702 | 39.90552 | 2.777466 | 0.525212 | 5.288273 | 1.23E-07 | 5.80E-05 | Gm6553 |
| ENSMUSG00000052387 | 2491.318 | 2.731205 | 0.523106 | 5.221127 | 1.78E-07 | 7.51E-05 | Trpm3 |
| ENSMUSG00000027674 | 986.0046 | 2.706418 | 0.487322 | 5.55365 | 2.80E-08 | 2.06E-05 | Pex5l |
| ENSMUSG00000070473 | 80.87005 | 2.70426 | 0.784546 | 3.446911 | 0.000567 | 0.033478 | Cldn3 |
| ENSMUSG00000071311 | 46.71959 | 2.697919 | 0.50327 | 5.360781 | 8.29E-08 | 4.64E-05 | Gpr31b |
| ENSMUSG00000019232 | 711.2605 | 2.676632 | 0.325869 | 8.213828 | 2.14E-16 | 2.13E-12 | Etnppl |
| ENSMUSG00000034839 | 281.8842 | 2.669664 | 0.640044 | 4.171065 | 3.03E-05 | 0.003855 | Larp6 |
| ENSMUSG00000025597 | 452.2456 | 2.638733 | 0.542415 | 4.864789 | 1.15E-06 | 0.000299 | Klhl4 |
| ENSMUSG00000056966 | 1134.402 | 2.626012 | 0.554571 | 4.735212 | 2.19E-06 | 0.000505 | Gjc3 |
| ENSMUSG00000019997 | 1001.017 | 2.578914 | 0.623618 | 4.135408 | 3.54E-05 | 0.004339 | Ccn2 |
| ENSMUSG00000029095 | 360.8656 | 2.578911 | 0.612218 | 4.212408 | 2.53E-05 | 0.003364 | Ablim2 |
| ENSMUSG00000095334 | 48.88959 | 2.576161 | 0.587445 | 4.385369 | 1.16E-05 | 0.001982 | NA |
| ENSMUSG00000040264 | 48.46615 | 2.572104 | 0.735399 | 3.497564 | 0.00047 | 0.029382 | Gbp2b |
| ENSMUSG00000050854 | 217.9323 | 2.56138 | 0.562885 | 4.550451 | 5.35E-06 | 0.001062 | Tmem125 |
| ENSMUSG00000015652 | 755.9357 | 2.555172 | 0.47683 | 5.358663 | 8.38E-08 | 4.64E-05 | Steap1 |
| ENSMUSG00000026830 | 1687.137 | 2.545722 | 0.476444 | 5.343172 | 9.13E-08 | 4.90E-05 | Ermn |
| ENSMUSG00000020486 | 4482.534 | 2.48817 | 0.478878 | 5.195831 | 2.04E-07 | 8.19E-05 | Septin4 |
| ENSMUSG00000044519 | 34.39078 | 2.487324 | 0.733883 | 3.389266 | 0.000701 | 0.037983 | Zfp488 |
| ENSMUSG00000073680 | 1508.183 | 2.479955 | 0.533514 | 4.648336 | 3.35E-06 | 0.000722 | Tmem88b |
| ENSMUSG00000021838 | 337.944 | 2.470195 | 0.640516 | 3.85657 | 0.000115 | 0.010811 | Samd4 |
| ENSMUSG00000027375 | 5023.143 | 2.460394 | 0.495106 | 4.969432 | 6.71E-07 | 0.000194 | Mal |
| ENSMUSG00000036098 | 613.7566 | 2.439961 | 0.473938 | 5.148271 | 2.63E-07 | 9.31E-05 | Myrf |
| ENSMUSG00000102099 | 57.86985 | 2.427682 | 0.48546 | 5.000784 | 5.71E-07 | 0.000177 | 1700011B04Rik |
| ENSMUSG00000037996 | 139.8738 | 2.404463 | 0.50626 | 4.749459 | 2.04E-06 | 0.000482 | Slc24a2 |
| ENSMUSG00000032988 | 507.9495 | 2.385277 | 0.562621 | 4.239579 | 2.24E-05 | 0.003043 | Slc16a8 |
| ENSMUSG00000116935 | 27.45575 | 2.37746 | 0.580852 | 4.093055 | 4.26E-05 | 0.004997 | Gpr31a |
| ENSMUSG00000091712 | 126.2919 | 2.345941 | 0.563135 | 4.165858 | 3.10E-05 | 0.003919 | Sec14l5 |
| ENSMUSG00000081225 | 91.59152 | 2.34501 | 0.615598 | 3.809318 | 0.000139 | 0.012231 | Cyp2j12 |
| ENSMUSG00000068323 | 728.2826 | 2.295528 | 0.457271 | 5.02006 | 5.17E-07 | 0.000168 | Slc4a5 |
| ENSMUSG00000015806 | 8009.753 | 2.262004 | 0.555072 | 4.075155 | 4.60E-05 | 0.005242 | Qdpr |
| ENSMUSG00000004864 | 88.61411 | 2.258971 | 0.654966 | 3.448988 | 0.000563 | 0.033419 | Mapk13 |
| ENSMUSG00000050121 | 975.2839 | 2.251817 | 0.520462 | 4.326574 | 1.51E-05 | 0.002366 | Opalin |
| ENSMUSG00000031425 | 38648.99 | 2.250395 | 0.411484 | 5.468969 | 4.53E-08 | 3.10E-05 | Plp1 |
| ENSMUSG00000037754 | 1073.881 | 2.232254 | 0.544812 | 4.097294 | 4.18E-05 | 0.004936 | Ppp1r16b |
| ENSMUSG00000057969 | 1008.143 | 2.211946 | 0.509872 | 4.338236 | 1.44E-05 | 0.002261 | Sema3b |
| ENSMUSG00000005125 | 2965.183 | 2.210331 | 0.583947 | 3.785158 | 0.000154 | 0.013119 | Ndrg1 |
| ENSMUSG00000039481 | 68.60899 | 2.210327 | 0.624235 | 3.540859 | 0.000399 | 0.02646 | Nrtn |
| ENSMUSG00000047797 | 280.6806 | 2.201973 | 0.518452 | 4.24721 | 2.16E-05 | 0.002982 | Gjb1 |
| ENSMUSG00000036745 | 958.1891 | 2.198588 | 0.591675 | 3.715869 | 0.000203 | 0.015972 | Ttll7 |
| ENSMUSG00000026686 | 162.4603 | 2.197829 | 0.622787 | 3.52902 | 0.000417 | 0.027217 | Lmx1a |

| | | | | | | | |
|---|---|---|---|---|---|---|---|
| ENSMUSG00000079436 | 1050.059 | 1.84974 | 0.45595 | 4.05689 | 4.97E-05 | 0.005511 | Kcnj13 |
| ENSMUSG00000025350 | 891.1418 | 1.845791 | 0.339351 | 5.439185 | 5.35E-08 | 3.54E-05 | Rdh5 |
| ENSMUSG00000026991 | 1092.899 | 1.843931 | 0.44792 | 4.116654 | 3.84E-05 | 0.004622 | Pkp4 |
| ENSMUSG00000037490 | 1313.525 | 1.827113 | 0.399888 | 4.56906 | 4.90E-06 | 0.000992 | Slc2a12 |
| ENSMUSG00000055235 | 361.4679 | 1.807402 | 0.420087 | 4.302449 | 1.69E-05 | 0.002558 | Wdr86 |
| ENSMUSG00000109325 | 76.95489 | 1.803811 | 0.528371 | 3.413908 | 0.00064 | 0.035484 | NA |
| ENSMUSG00000021097 | 578.4045 | 1.798505 | 0.30171 | 5.961039 | 2.51E-09 | 2.92E-06 | Clmn |
| ENSMUSG00000003934 | 1481.149 | 1.795616 | 0.409477 | 4.385142 | 1.16E-05 | 0.001982 | Efnb3 |
| ENSMUSG00000002769 | 118.7657 | 1.781723 | 0.529309 | 3.366131 | 0.000762 | 0.040166 | Gnmt |
| ENSMUSG00000035189 | 158.3332 | 1.775346 | 0.459619 | 3.862646 | 0.000112 | 0.010595 | Ano4 |
| ENSMUSG00000026109 | 726.303 | 1.774024 | 0.378803 | 4.68324 | 2.82E-06 | 0.000622 | Tmeff2 |
| ENSMUSG00000039672 | 691.6759 | 1.764924 | 0.479427 | 3.68132 | 0.000232 | 0.017703 | Kcne2 |
| ENSMUSG00000023043 | 810.8047 | 1.763177 | 0.420442 | 4.193625 | 2.75E-05 | 0.003559 | Krt18 |
| ENSMUSG00000057286 | 389.3667 | 1.750603 | 0.484396 | 3.613988 | 0.000302 | 0.021438 | St6galnac2 |
| ENSMUSG00000038526 | 842.3208 | 1.750046 | 0.418285 | 4.183861 | 2.87E-05 | 0.003692 | Car14 |
| ENSMUSG00000041380 | 286.4886 | 1.747209 | 0.501728 | 3.482385 | 0.000497 | 0.030904 | Htr2c |
| ENSMUSG00000020108 | 2104.028 | 1.744534 | 0.329059 | 5.301577 | 1.15E-07 | 5.61E-05 | Ddit4 |
| ENSMUSG00000021662 | 343.3016 | 1.733994 | 0.434665 | 3.989263 | 6.63E-05 | 0.006993 | Arhgef28 |
| ENSMUSG00000049044 | 3055.881 | 1.732367 | 0.463991 | 3.733622 | 0.000189 | 0.015158 | Rapgef4 |
| ENSMUSG00000026437 | 196.4548 | 1.718317 | 0.318558 | 5.394048 | 6.89E-08 | 4.41E-05 | Cdk18 |
| ENSMUSG00000000792 | 134.627 | 1.708788 | 0.412208 | 4.145449 | 3.39E-05 | 0.004179 | Slc5a5 |
| ENSMUSG00000033006 | 478.3561 | 1.701545 | 0.51513 | 3.303135 | 0.000956 | 0.04767 | Sox10 |
| ENSMUSG00000046160 | 2037.975 | 1.692992 | 0.396364 | 4.271306 | 1.94E-05 | 0.002785 | Olig1 |
| ENSMUSG00000039809 | 444.2229 | 1.691937 | 0.512455 | 3.301632 | 0.000961 | 0.04767 | Gabbr2 |
| ENSMUSG00000022472 | 2939.752 | 1.683604 | 0.419604 | 4.01236 | 6.01E-05 | 0.006481 | Desi1 |
| ENSMUSG00000039904 | 1927.182 | 1.675534 | 0.35199 | 4.760179 | 1.93E-06 | 0.000468 | Gpr37 |
| ENSMUSG00000041801 | 546.1416 | 1.661447 | 0.467624 | 3.552957 | 0.000381 | 0.025615 | Phlda3 |
| ENSMUSG00000031605 | 1790.779 | 1.658285 | 0.426797 | 3.885418 | 0.000102 | 0.010082 | Klhl2 |
| ENSMUSG00000030616 | 306.8899 | 1.657122 | 0.468624 | 3.536146 | 0.000406 | 0.026758 | Sytl2 |
| ENSMUSG00000074578 | 512.729 | 1.654372 | 0.178362 | 9.275377 | 1.77E-20 | 3.51E-16 | Zfas1 |
| ENSMUSG00000049382 | 366.923 | 1.644069 | 0.4751 | 3.46047 | 0.000539 | 0.032513 | Krt8 |
| ENSMUSG00000015970 | 163.1632 | 1.641638 | 0.448891 | 3.657097 | 0.000255 | 0.018811 | Chdh |
| ENSMUSG00000049811 | 258.9241 | 1.640745 | 0.464218 | 3.534431 | 0.000409 | 0.026843 | Fam161a |
| ENSMUSG00000079597 | 87.78939 | 1.633071 | 0.343517 | 4.753981 | 1.99E-06 | 0.000477 | Cstdc4 |
| ENSMUSG00000041205 | 522.3426 | 1.633013 | 0.431515 | 3.784372 | 0.000154 | 0.013119 | Map6d1 |
| ENSMUSG00000095620 | 43.94096 | 1.623875 | 0.443419 | 3.662166 | 0.00025 | 0.018581 | Csta2 |
| ENSMUSG00000031465 | 242.6927 | 1.6211 | 0.437682 | 3.70383 | 0.000212 | 0.016456 | Angpt2 |
| ENSMUSG00000042686 | 157.8817 | 1.619504 | 0.423539 | 3.823745 | 0.000131 | 0.011681 | Jph1 |
| ENSMUSG00000031444 | 387.1771 | 1.618053 | 0.397924 | 4.066237 | 4.78E-05 | 0.005385 | F10 |
| ENSMUSG00000036634 | 5180.545 | 1.617443 | 0.422917 | 3.824494 | 0.000131 | 0.011681 | Mag |
| ENSMUSG00000086290 | 104.8313 | 1.610706 | 0.3705 | 4.347388 | 1.38E-05 | 0.002204 | Snhg12 |
| ENSMUSG00000074170 | 247.6764 | 1.608936 | 0.488439 | 3.294039 | 0.000988 | 0.048855 | Plekhf1 |
| ENSMUSG00000021187 | 90.66386 | 1.60274 | 0.414182 | 3.86965 | 0.000109 | 0.010445 | Tc2n |
| ENSMUSG00000049176 | 245.0562 | 1.598331 | 0.325367 | 4.912397 | 9.00E-07 | 0.000255 | Frmpd4 |
| ENSMUSG00000032060 | 2557.833 | 1.570832 | 0.423031 | 3.713276 | 0.000205 | 0.016042 | Cryab |
| ENSMUSG00000074093 | 1511.593 | 1.561873 | 0.46363 | 3.368789 | 0.000755 | 0.039938 | Svip |
| ENSMUSG00000029816 | 77.16312 | 1.554747 | 0.442916 | 3.510254 | 0.000448 | 0.028412 | Gpnmb |

| | | | | | | | |
|---|---|---|---|---|---|---|---|
| ENSMUSG00000049493 | 405.6949 | 1.546699 | 0.449419 | 3.441552 | 0.000578 | 0.033745 | Pls1 |
| ENSMUSG00000092536 | 121.0009 | 1.515374 | 0.451253 | 3.358143 | 0.000785 | 0.04107 | NA |
| ENSMUSG00000006782 | 17261.55 | 1.490601 | 0.402983 | 3.698917 | 0.000217 | 0.016648 | Cnp |
| ENSMUSG00000025488 | 227.8406 | 1.489232 | 0.432103 | 3.446473 | 0.000568 | 0.033478 | Cox8b |
| ENSMUSG00000042429 | 825.1056 | 1.462245 | 0.302901 | 4.827468 | 1.38E-06 | 0.000352 | Adora1 |
| ENSMUSG00000022425 | 74438.3 | 1.457523 | 0.420986 | 3.462163 | 0.000536 | 0.032408 | Enpp2 |
| ENSMUSG00000040473 | 252.3865 | 1.444804 | 0.271181 | 5.327822 | 9.94E-08 | 5.19E-05 | Cfap69 |
| ENSMUSG00000062591 | 7270.1 | 1.441612 | 0.379165 | 3.802071 | 0.000143 | 0.012338 | Tubb4a |
| ENSMUSG00000028328 | 462.0067 | 1.438158 | 0.25427 | 5.656036 | 1.55E-08 | 1.28E-05 | Tmod1 |
| ENSMUSG00000032854 | 832.2016 | 1.434627 | 0.414748 | 3.459036 | 0.000542 | 0.032588 | Ugt8a |
| ENSMUSG00000021848 | 484.1138 | 1.430915 | 0.332135 | 4.308237 | 1.65E-05 | 0.002531 | Otx2 |
| ENSMUSG00000053475 | 727.0577 | 1.420798 | 0.415293 | 3.421192 | 0.000623 | 0.035216 | Tnfaip6 |
| ENSMUSG00000044734 | 447.2619 | 1.417877 | 0.424674 | 3.338746 | 0.000842 | 0.043362 | Serpinb1a |
| ENSMUSG00000030142 | 1186.544 | 1.401086 | 0.268636 | 5.215552 | 1.83E-07 | 7.57E-05 | Clec4e |
| ENSMUSG00000115625 | 343.3371 | 1.399546 | 0.374832 | 3.733798 | 0.000189 | 0.015158 | 2900040C04Rik |
| ENSMUSG00000049907 | 326.2262 | 1.392803 | 0.408714 | 3.407766 | 0.000655 | 0.035891 | Rasl11b |
| ENSMUSG00000079470 | 319.8967 | 1.386504 | 0.329579 | 4.206892 | 2.59E-05 | 0.003416 | Utp14b |
| ENSMUSG00000050914 | 156.9798 | 1.378534 | 0.374705 | 3.678987 | 0.000234 | 0.017797 | Ankrd37 |
| ENSMUSG00000040260 | 521.9889 | 1.374305 | 0.327287 | 4.199089 | 2.68E-05 | 0.003497 | Daam2 |
| ENSMUSG00000038077 | 323.9781 | 1.370848 | 0.344045 | 3.984502 | 6.76E-05 | 0.007097 | Kcna6 |
| ENSMUSG00000022949 | 2514.072 | 1.36482 | 0.367265 | 3.71617 | 0.000202 | 0.015972 | Clic6 |
| ENSMUSG00000048498 | 1515.198 | 1.345709 | 0.28034 | 4.800271 | 1.58E-06 | 0.000398 | Cd300e |
| ENSMUSG00000042662 | 81.96791 | 1.337976 | 0.398649 | 3.356277 | 0.00079 | 0.041131 | Dusp15 |
| ENSMUSG00000031431 | 13044.41 | 1.324317 | 0.313083 | 4.229918 | 2.34E-05 | 0.003155 | Tsc22d3 |
| ENSMUSG00000003849 | 548.2518 | 1.310647 | 0.250506 | 5.231994 | 1.68E-07 | 7.23E-05 | Nqo1 |
| ENSMUSG00000051748 | 267.5511 | 1.305821 | 0.258472 | 5.052078 | 4.37E-07 | 0.000145 | Wfdc21 |
| ENSMUSG00000027570 | 2597.201 | 1.303894 | 0.313652 | 4.15713 | 3.22E-05 | 0.004021 | Col9a3 |
| ENSMUSG00000020681 | 3448.194 | 1.295233 | 0.333269 | 3.886448 | 0.000102 | 0.010082 | Ace |
| ENSMUSG00000045551 | 759.653 | 1.294704 | 0.290835 | 4.451687 | 8.52E-06 | 0.00158 | Fpr1 |
| ENSMUSG00000033208 | 1249.084 | 1.285337 | 0.327145 | 3.928956 | 8.53E-05 | 0.008591 | S100b |
| ENSMUSG00000027562 | 7311.703 | 1.276394 | 0.242415 | 5.265323 | 1.40E-07 | 6.31E-05 | Car2 |
| ENSMUSG00000027030 | 1674.525 | 1.247736 | 0.347446 | 3.591162 | 0.000329 | 0.022995 | Stk39 |
| ENSMUSG00000047617 | 1230.892 | 1.247631 | 0.270563 | 4.611237 | 4.00E-06 | 0.000836 | Paxx |
| ENSMUSG00000036585 | 1140.249 | 1.239959 | 0.356998 | 3.473299 | 0.000514 | 0.031672 | Fgf1 |
| ENSMUSG00000032246 | 621.3906 | 1.234425 | 0.360193 | 3.42712 | 0.00061 | 0.034873 | Calml4 |
| ENSMUSG00000037664 | 1116.614 | 1.222372 | 0.326256 | 3.746665 | 0.000179 | 0.014703 | Cdkn1c |
| ENSMUSG00000032579 | 898.3502 | 1.219534 | 0.30059 | 4.057135 | 4.97E-05 | 0.005511 | Hemk1 |
| ENSMUSG00000055137 | 190.0163 | 1.210709 | 0.303143 | 3.99386 | 6.50E-05 | 0.006896 | Sugct |
| ENSMUSG00000014791 | 115.1717 | 1.208739 | 0.339584 | 3.559465 | 0.000372 | 0.025272 | Elmo3 |
| ENSMUSG00000048022 | 1972.043 | 1.189737 | 0.306974 | 3.875691 | 0.000106 | 0.010286 | Tmem229a |
| ENSMUSG00000035112 | 150.4751 | 1.178173 | 0.326448 | 3.609073 | 0.000307 | 0.021771 | Wnk4 |
| ENSMUSG00000029053 | 892.5281 | 1.171757 | 0.341644 | 3.429755 | 0.000604 | 0.034636 | Prkcz |
| ENSMUSG00000072596 | 1520.173 | 1.166051 | 0.290997 | 4.007094 | 6.15E-05 | 0.006591 | Ear2 |
| ENSMUSG00000110030 | 162.7707 | 1.162193 | 0.335286 | 3.466273 | 0.000528 | 0.032159 | NA |
| ENSMUSG00000025586 | 185.6149 | 1.151753 | 0.270715 | 4.254488 | 2.10E-05 | 0.002915 | Cpeb1 |
| ENSMUSG00000033102 | 216.9797 | 1.145381 | 0.335065 | 3.418385 | 0.00063 | 0.035216 | Cdc14b |
| ENSMUSG00000090166 | 268.6031 | 1.138957 | 0.335423 | 3.395583 | 0.000685 | 0.037321 | Ear10 |

| | | | | | | | |
|---|---|---|---|---|---|---|---|
| ENSMUSG00000023827 | 1894.244 | 1.138073 | 0.326981 | 3.480547 | 0.0005 | 0.030976 | Agpat4 |
| ENSMUSG00000019856 | 287.4046 | 1.135891 | 0.317594 | 3.576544 | 0.000348 | 0.024065 | Fam184a |
| ENSMUSG00000006931 | 747.1615 | 1.133698 | 0.321087 | 3.530815 | 0.000414 | 0.027122 | P3h4 |
| ENSMUSG00000063458 | 141.8336 | 1.132071 | 0.342117 | 3.309018 | 0.000936 | 0.046899 | Lrmda |
| ENSMUSG00000030711 | 2560.403 | 1.131901 | 0.211236 | 5.358476 | 8.39E-08 | 4.64E-05 | Sult1a1 |
| ENSMUSG00000022974 | 1406.676 | 1.126112 | 0.319902 | 3.520176 | 0.000431 | 0.027866 | Paxbp1 |
| ENSMUSG00000086841 | 766.0955 | 1.12277 | 0.222243 | 5.052001 | 4.37E-07 | 0.000145 | 2410006H16Rik |
| ENSMUSG00000030087 | 1524.984 | 1.11836 | 0.228955 | 4.884636 | 1.04E-06 | 0.000285 | Klf15 |
| ENSMUSG00000043822 | 146.0396 | 1.117313 | 0.300289 | 3.720791 | 0.000199 | 0.015886 | Adamtsl5 |
| ENSMUSG00000044033 | 675.2554 | 1.110127 | 0.332824 | 3.335477 | 0.000852 | 0.043761 | Ccdc141 |
| ENSMUSG00000026203 | 2212.328 | 1.090434 | 0.314769 | 3.464233 | 0.000532 | 0.032258 | Dnajb2 |
| ENSMUSG00000019577 | 820.8845 | 1.086944 | 0.267154 | 4.068596 | 4.73E-05 | 0.005361 | Pdk4 |
| ENSMUSG00000066682 | 712.0235 | 1.079304 | 0.319704 | 3.375944 | 0.000736 | 0.039228 | Pilrb2 |
| ENSMUSG00000053332 | 2619.585 | 1.074093 | 0.214799 | 5.000467 | 5.72E-07 | 0.000177 | Gas5 |
| ENSMUSG00000033039 | 758.1241 | 1.068986 | 0.277716 | 3.84921 | 0.000118 | 0.010984 | Micall1 |
| ENSMUSG00000005800 | 613.5976 | 1.067186 | 0.321833 | 3.315959 | 0.000913 | 0.046099 | Mmp8 |
| ENSMUSG00000003526 | 1577.611 | 1.066883 | 0.248267 | 4.297321 | 1.73E-05 | 0.002578 | Prodh |
| ENSMUSG00000091586 | 187.1814 | 1.055915 | 0.300333 | 3.515815 | 0.000438 | 0.028145 | Cyp4f17 |
| ENSMUSG00000113029 | 107.872 | 1.052933 | 0.308739 | 3.410429 | 0.000649 | 0.03574 | NA |
| ENSMUSG00000024222 | 4655.745 | 1.051946 | 0.170318 | 6.176364 | 6.56E-10 | 1.06E-06 | Fkbp5 |
| ENSMUSG00000031791 | 699.4264 | 1.021442 | 0.256656 | 3.979812 | 6.90E-05 | 0.007134 | Tmem38a |
| ENSMUSG00000038550 | 273.5747 | 1.011528 | 0.294826 | 3.430932 | 0.000602 | 0.034586 | Ciart |
| ENSMUSG00000006651 | 15002.21 | 1.002897 | 0.27583 | 3.635925 | 0.000277 | 0.020053 | Aplp1 |
| ENSMUSG00000039470 | 330.7039 | -1.00672 | 0.24855 | -4.05036 | 5.11E-05 | 0.005605 | Zdhhc2 |
| ENSMUSG00000025001 | 546.3061 | -1.01227 | 0.25418 | -3.98251 | 6.82E-05 | 0.00712 | Hells |
| ENSMUSG00000027848 | 59939.07 | -1.04913 | 0.188107 | -5.57729 | 2.44E-08 | 1.86E-05 | Olfml3 |
| ENSMUSG00000025665 | 422.0151 | -1.06175 | 0.263877 | -4.02364 | 5.73E-05 | 0.006212 | Rps6ka6 |
| ENSMUSG00000090122 | 727.0424 | -1.12873 | 0.197013 | -5.72924 | 1.01E-08 | 9.10E-06 | Kcne1l |
| ENSMUSG00000027350 | 333.3992 | -1.15226 | 0.32845 | -3.50819 | 0.000451 | 0.028412 | Chgb |
| ENSMUSG00000079445 | 105.4999 | -1.15487 | 0.311665 | -3.70549 | 0.000211 | 0.016413 | B3gnt7 |
| ENSMUSG00000072082 | 763.2923 | -1.2126 | 0.359178 | -3.37604 | 0.000735 | 0.039228 | Ccnf |
| ENSMUSG00000021194 | 389.0526 | -1.3547 | 0.302943 | -4.47181 | 7.76E-06 | 0.001465 | Chga |
| ENSMUSG00000060780 | 129.244 | -1.40977 | 0.406756 | -3.46588 | 0.000528 | 0.032159 | Lrrtm1 |
| ENSMUSG00000034656 | 248.7135 | -1.44962 | 0.337269 | -4.29811 | 1.72E-05 | 0.002578 | Cacna1a |
| ENSMUSG00000000037 | 88.53711 | -1.45658 | 0.426153 | -3.41798 | 0.000631 | 0.035216 | Scml2 |
| ENSMUSG00000021356 | 166.7265 | -1.48858 | 0.441812 | -3.36927 | 0.000754 | 0.039938 | Irf4 |
| ENSMUSG00000004612 | 86.66066 | -1.51003 | 0.429459 | -3.51613 | 0.000438 | 0.028145 | Nkg7 |
| ENSMUSG00000073643 | 2946.989 | -1.57658 | 0.329021 | -4.79171 | 1.65E-06 | 0.00041 | Wdfy1 |
| ENSMUSG00000061186 | 97.76425 | -1.72365 | 0.422278 | -4.0818 | 4.47E-05 | 0.005154 | Sfmbt2 |
| ENSMUSG00000046204 | 305.3188 | -1.7292 | 0.380413 | -4.54559 | 5.48E-06 | 0.001076 | Pnma2 |
| ENSMUSG00000009350 | 39.73776 | -4.84473 | 0.826571 | -5.86124 | 4.59E-09 | 4.56E-06 | Mpo |
| ENSMUSG00000076514 | 21.86559 | -5.42295 | 1.33045 | -4.07602 | 4.58E-05 | 0.005242 | Igkv17-121 |
| ENSMUSG00000071636 | 25.35944 | -7.14267 | 1.511443 | -4.72573 | 2.29E-06 | 0.000523 | Rimbp3 |
