## Supplemental Table S2 for "Astrocytic Nonsense-mediated mRNA decay regulates calcium signaling to support synapse function and influence anxiety-like behavior"

**Table S2. NMD inducing feature analysis of DEGs**

| Transcript_id | dEJ | uORF | 3'UTR | |
|---|---|---|---|---|
| - | - | - | - | |
| ENSMUST00000039450 | FALSE | FALSE | 1208 | Mcoln3 |
| ENSMUST00000027639 | FALSE | FALSE | 121 | Marco |
| ENSMUST00000100926 | FALSE | FALSE | 156 | Tex52 |
| ENSMUST00000038730 | FALSE | FALSE | 2282 | Rtp1 |
| ENSMUST00000068698 | FALSE | FALSE | 849 | Mobp |
| ENSMUST00000103328 | 0 | FALSE | 0 | Igkv10-96 |
| ENSMUST00000000808 | FALSE | FALSE | 553 | Il12rb1 |
| ENSMUST00000006956 | FALSE | FALSE | 128 | Saa3 |
| ENSMUST00000047865 | FALSE | FALSE | 1481 | Mbp |
| ENSMUST00000006969 | FALSE | FALSE | 223 | Krt23 |
| ENSMUST00000013667 | FALSE | FALSE | 926 | Bcas1 |
| - | - | - | - | |
| ENSMUST00000024738 | FALSE | TRUE | 457 | Sult1c1 |
| ENSMUST00000061469 | FALSE | FALSE | 1642 | Wfikkn2 |
| - | - | - | - | |
| ENSMUST00000044002 | FALSE | FALSE | 530 | Tmem184a |
| ENSMUST00000099362 | FALSE | FALSE | 321 | Slc4a11 |
| ENSMUST00000050962 | FALSE | TRUE | 205 | Cldn14 |
| ENSMUST00000069742 | FALSE | FALSE | 2176 | Prr18 |
| ENSMUST00000037434 | FALSE | FALSE | 2272 | Cacna2d4 |
| ENSMUST00000017878 | FALSE | TRUE | 432 | Dbndd2 |
| - | - | - | - | |
| ENSMUST00000002663 | FALSE | TRUE | 248 | Pon1 |
| ENSMUST00000023886 | FALSE | TRUE | 365 | Sult1c2 |
| ENSMUST00000045902 | FALSE | FALSE | 2411 | Fmo2 |
| ENSMUST00000015545 | FALSE | FALSE | 507 | Cltrn |
| ENSMUST00000079176 | FALSE | FALSE | 1034 | Plekhb1 |
| ENSMUST00000104937 | 0 | FALSE | 3163 | Ankrd63 |
| - | - | - | - | |
| ENSMUST00000066532 | FALSE | FALSE | 1995 | Lipg |
| - | - | - | - | |
| ENSMUST00000026573 | FALSE | FALSE | 81 | Lmntd2 |
| - | - | - | - | |
| ENSMUST00000032874 | FALSE | FALSE | 424 | Sh3gl3 |
| ENSMUST00000044792 | FALSE | TRUE | 2383 | Rcan2 |
| ENSMUST00000005014 | FALSE | FALSE | 481 | Hapln2 |
| ENSMUST00000096089 | FALSE | FALSE | 78 | Cstdc5 |
| ENSMUST00000020702 | FALSE | FALSE | 1435 | Igfbp3 |
| ENSMUST00000042097 | FALSE | FALSE | 78 | Stfa1 |
| ENSMUST00000114850 | FALSE | FALSE | 57 | Cstdc6 |
| - | - | - | - | |
| - | - | - | - | |
| - | - | - | - | |
| ENSMUST00000097974 | TRUE | FALSE | 815 | Nkx6-2 |
| ENSMUST00000023619 | FALSE | FALSE | 35 | Stfa2 |

Orange: upregulated genes
Green: downregulated genes
yellow: contain at least one NMD feature

| | | | | |
|---|---|---|---|---|
| ENSMUST00000055018 | FALSE | FALSE | 2078 | Lpar1 |
| - | - | - | - | |
| ENSMUST00000132795 | 0 | TRUE | 0 | Tnni1 |
| ENSMUST00000002908 | FALSE | FALSE | 1416 | Plin4 |
| - | - | - | - | |
| - | - | - | - | |
| ENSMUST00000052458 | FALSE | FALSE | 1015 | Lrrc38 |
| - | - | - | - | |
| ENSMUST00000037901 | FALSE | FALSE | 494 | Trpm3 |
| ENSMUST00000078226 | FALSE | FALSE | 1035 | Pex5l |
| ENSMUST00000094245 | 0 | FALSE | 368 | Cldn3 |
| ENSMUST00000091648 | 0 | FALSE | 307 | Gpr31b |
| ENSMUST00000072271 | FALSE | FALSE | 2844 | Etnppl |
| ENSMUST00000038407 | FALSE | FALSE | 744 | Larp6 |
| ENSMUST00000040504 | FALSE | FALSE | 1339 | Klhl4 |
| ENSMUST00000077119 | FALSE | FALSE | 2991 | Gjc3 |
| ENSMUST00000020171 | FALSE | FALSE | 1067 | Ccn2 |
| ENSMUST00000054598 | FALSE | FALSE | 1554 | Ablim2 |
| - | - | - | - | |
| ENSMUST00000029936 | FALSE | FALSE | 974 | Gbp2b |
| ENSMUST00000060214 | FALSE | TRUE | 641 | Tmem125 |
| ENSMUST00000015796 | FALSE | FALSE | 99 | Steap1 |
| ENSMUST00000090940 | FALSE | FALSE | 2641 | Ermn |
| ENSMUST00000018544 | FALSE | FALSE | 164 | Septin4 |
| ENSMUST00000166737 | FALSE | FALSE | 3121 | Zfp488 |
| ENSMUST00000097742 | FALSE | FALSE | 2677 | Tmem88b |
| ENSMUST00000022386 | FALSE | TRUE | 1691 | Samd4 |
| ENSMUST00000028853 | FALSE | FALSE | 106 | Mal |
| ENSMUST00000088013 | FALSE | FALSE | 2113 | Myrf |
| - | - | - | - | |
| ENSMUST00000044990 | FALSE | FALSE | 171 | Slc24a2 |
| ENSMUST00000039752 | FALSE | FALSE | 114 | Slc16a8 |
| - | - | - | - | |
| ENSMUST00000165810 | FALSE | FALSE | 184 | Sec14l5 |
| ENSMUST00000097972 | FALSE | FALSE | 70 | Cyp2j12 |
| ENSMUST00000039212 | FALSE | FALSE | 1652 | Slc4a5 |
| ENSMUST00000015950 | FALSE | FALSE | 520 | Qdpr |
| ENSMUST00000004986 | FALSE | 0 | 599 | Mapk13 |
| ENSMUST00000087176 | FALSE | FALSE | 1330 | Opalin |
| ENSMUST00000033800 | FALSE | FALSE | 3622 | Plp1 |
| ENSMUST00000045503 | FALSE | TRUE | 4471 | Ppp1r16b |
| ENSMUST00000073448 | FALSE | TRUE | 417 | Sema3b |
| ENSMUST00000005256 | FALSE | FALSE | 1549 | Ndrg1 |
| ENSMUST00000044752 | FALSE | TRUE | 87 | Nrtn |
| ENSMUST00000052130 | FALSE | FALSE | 610 | Gjb1 |
| ENSMUST00000037942 | FALSE | FALSE | 4258 | Ttll7 |
| ENSMUST00000028003 | FALSE | FALSE | 1980 | Lmx1a |

| | | | | |
|---|---|---|---|---|
| ENSMUST00000033169 | FALSE | FALSE | 897 | Cdr2 |
| ENSMUST00000029451 | FALSE | TRUE | 3350 | Tspan2 |
| ENSMUST00000036631 | FALSE | TRUE | 659 | Dusp26 |
| ENSMUST00000046174 | FALSE | FALSE | 1016 | Cldn11 |
| - | - | - | - | |
| ENSMUST00000039928 | FALSE | FALSE | 2062 | Plekhh1 |
| ENSMUST00000070139 | FALSE | FALSE | 1079 | Cndp1 |
| ENSMUST00000015664 | FALSE | FALSE | 440 | Ctsk |
| ENSMUST00000045224 | FALSE | FALSE | 3295 | Acer2 |
| ENSMUST00000075312 | FALSE | FALSE | 583 | Ttr |
| ENSMUST00000185502 | FALSE | FALSE | 3201 | Spock1 |
| ENSMUST00000077066 | FALSE | FALSE | 2440 | Tmem151a |
| ENSMUST00000102665 | FALSE | TRUE | 761 | Mog |
| ENSMUST00000004774 | FALSE | FALSE | 1758 | Aqp1 |
| ENSMUST00000054484 | FALSE | FALSE | 1939 | Slc4a10 |
| ENSMUST00000127305 | FALSE | TRUE | 1552 | Epn3 |
| ENSMUST00000063004 | FALSE | TRUE | 249 | Gal3st1 |
| - | - | - | - | |
| ENSMUST00000079038 | FALSE | FALSE | 6480 | Hhip |
| ENSMUST00000106332 | FALSE | FALSE | 188 | Sez6l2 |
| ENSMUST00000190196 | 0 | FALSE | 1816 | Prob1 |
| - | - | - | - | |
| ENSMUST00000032633 | FALSE | FALSE | 489 | Oca2 |
| ENSMUST00000064433 | FALSE | FALSE | 8465 | Tmod2 |
| ENSMUST00000106770 | FALSE | TRUE | 221 | Mroh7 |
| - | - | - | - | |
| - | - | - | - | |
| ENSMUST00000013995 | FALSE | FALSE | 226 | Abca4 |
| ENSMUST00000069330 | FALSE | FALSE | 293 | Acp5 |
| ENSMUST00000015234 | FALSE | FALSE | 162 | Ptgds |
| - | - | - | - | |
| ENSMUST00000025563 | FALSE | FALSE | 156 | Fth1 |
| ENSMUST00000026672 | FALSE | TRUE | 1011 | Pde8a |
| ENSMUST00000031868 | FALSE | TRUE | 874 | Slc13a4 |
| ENSMUST00000164834 | 0 | TRUE | 689 | Gpr62 |
| ENSMUST00000085989 | 0 | TRUE | 412 | Cldn9 |
| ENSMUST00000025724 | FALSE | FALSE | 1663 | Il33 |
| ENSMUST00000038475 | FALSE | FALSE | 1301 | Fa2h |
| ENSMUST00000030581 | FALSE | FALSE | 331 | Azin2 |
| ENSMUST00000062108 | FALSE | TRUE | 708 | Ikbke |
| ENSMUST00000071889 | FALSE | FALSE | 2488 | Car12 |
| ENSMUST00000021119 | FALSE | FALSE | 444 | Aspa |
| ENSMUST00000025800 | FALSE | TRUE | 758 | Pip5k1b |
| ENSMUST00000015797 | FALSE | TRUE | 1918 | Steap2 |
| ENSMUST00000043237 | FALSE | FALSE | 3058 | Trp53inp2 |
| ENSMUST00000047331 | FALSE | FALSE | 1290 | Lgi3 |
| ENSMUST00000122088 | FALSE | FALSE | 1014 | S1pr5 |

| | | | | |
|---|---|---|---|---|
| ENSMUST00000113212 | FALSE | FALSE | 53 | Kcnj13 |
| ENSMUST00000026406 | FALSE | FALSE | 120 | Rdh5 |
| ENSMUST00000037903 | FALSE | FALSE | 760 | Pkp4 |
| ENSMUST00000042261 | FALSE | TRUE | 2110 | Slc2a12 |
| ENSMUST00000068693 | FALSE | TRUE | 837 | Wdr86 |
| - | - | - | - | |
| ENSMUST00000109936 | FALSE | FALSE | 8675 | Clmn |
| ENSMUST00000004036 | FALSE | FALSE | 1768 | Efnb3 |
| ENSMUST00000002846 | FALSE | FALSE | 136 | Gnmt |
| ENSMUST00000181976 | FALSE | TRUE | 364 | Ano4 |
| ENSMUST00000081851 | FALSE | TRUE | 1789 | Tmeff2 |
| ENSMUST00000047383 | FALSE | FALSE | 1171 | Kcne2 |
| ENSMUST00000023803 | FALSE | FALSE | 66 | Krt18 |
| ENSMUST00000079545 | FALSE | FALSE | 2211 | St6galnac2 |
| ENSMUST00000036181 | FALSE | FALSE | 378 | Car14 |
| ENSMUST00000036303 | FALSE | TRUE | 2679 | Htr2c |
| ENSMUST00000020308 | FALSE | FALSE | 867 | Ddit4 |
| ENSMUST00000109426 | FALSE | FALSE | 178 | Arhgef28 |
| ENSMUST00000028525 | FALSE | FALSE | 1096 | Rapgef4 |
| ENSMUST00000027697 | FALSE | FALSE | 2699 | Cdk18 |
| ENSMUST00000000809 | FALSE | FALSE | 975 | Slc5a5 |
| ENSMUST00000040019 | FALSE | FALSE | 1025 | Sox10 |
| ENSMUST00000056882 | 0 | FALSE | 1274 | Olig1 |
| ENSMUST00000107749 | FALSE | FALSE | 2479 | Gabbr2 |
| ENSMUST00000023110 | TRUE | FALSE | 528 | Desi1 |
| ENSMUST00000054867 | FALSE | TRUE | 1538 | Gpr37 |
| ENSMUST00000038945 | TRUE | FALSE | 727 | Phlda3 |
| ENSMUST00000034017 | FALSE | TRUE | 1271 | Klhl2 |
| ENSMUST00000098310 | FALSE | FALSE | 891 | Sytl2 |
| - | - | - | - | |
| ENSMUST00000023952 | FALSE | FALSE | 242 | Krt8 |
| ENSMUST00000067620 | FALSE | FALSE | 3637 | Chdh |
| ENSMUST00000058269 | FALSE | TRUE | 184 | Fam161a |
| ENSMUST00000114858 | FALSE | FALSE | 72 | Cstdc4 |
| ENSMUST00000040880 | FALSE | FALSE | 2659 | Map6d1 |
| ENSMUST00000187183 | FALSE | FALSE | 72 | Csta2 |
| ENSMUST00000033846 | FALSE | TRUE | 1785 | Angpt2 |
| ENSMUST00000038382 | FALSE | FALSE | 2541 | Jph1 |
| ENSMUST00000033821 | FALSE | FALSE | 18 | F10 |
| ENSMUST00000040548 | FALSE | FALSE | 528 | Mag |
| - | - | - | - | |
| ENSMUST00000098513 | FALSE | FALSE | 4331 | Plekhf1 |
| ENSMUST00000110047 | TRUE | TRUE | 265 | Tc2n |
| ENSMUST00000112145 | FALSE | 0 | 3985 | Frmpd4 |
| ENSMUST00000034562 | FALSE | TRUE | 48 | Cryab |
| ENSMUST00000098414 | FALSE | FALSE | 2592 | Svip |
| ENSMUST00000031840 | FALSE | FALSE | 1868 | Gpnmb |

| | | | | |
|---|---|---|---|---|
| ENSMUST00000093800 | FALSE | TRUE | 1660 | Pls1 |
| - | - | - | - | |
| ENSMUST00000103120 | FALSE | TRUE | 932 | Cnp |
| ENSMUST00000026561 | FALSE | FALSE | 43 | Cox8b |
| ENSMUST00000038191 | FALSE | TRUE | 1602 | Adora1 |
| ENSMUST00000041591 | FALSE | FALSE | 465 | Enpp2 |
| ENSMUST00000054865 | FALSE | TRUE | 1918 | Cfap69 |
| ENSMUST00000071135 | FALSE | FALSE | 624 | Tubb4a |
| ENSMUST00000107773 | FALSE | FALSE | 1654 | Tmod1 |
| ENSMUST00000057944 | FALSE | FALSE | 1945 | Ugt8a |
| ENSMUST00000118578 | FALSE | FALSE | 1027 | Otx2 |
| ENSMUST00000065927 | FALSE | TRUE | 715 | Tnfaip6 |
| ENSMUST00000076352 | FALSE | FALSE | 726 | Serpinb1a |
| ENSMUST00000032239 | FALSE | FALSE | 1751 | Clec4e |
| - | - | - | - | |
| ENSMUST00000051937 | FALSE | FALSE | 902 | Rasl11b |
| ENSMUST00000053760 | FALSE | FALSE | 4783 | Utp14b |
| ENSMUST00000053558 | FALSE | FALSE | 223 | Ankrd37 |
| ENSMUST00000057610 | FALSE | FALSE | 99 | Daam2 |
| ENSMUST00000040751 | TRUE | FALSE | 3499 | Kcna6 |
| ENSMUST00000023670 | FALSE | FALSE | 1647 | Clic6 |
| ENSMUST00000062787 | FALSE | FALSE | 1344 | Cd300e |
| ENSMUST00000037715 | FALSE | FALSE | 170 | Dusp15 |
| ENSMUST00000055738 | FALSE | FALSE | 1340 | Tsc22d3 |
| ENSMUST00000003947 | FALSE | FALSE | 594 | Nqo1 |
| ENSMUST00000070832 | FALSE | FALSE | 155 | Wfdc21 |
| ENSMUST00000103059 | FALSE | FALSE | 765 | Col9a3 |
| ENSMUST00000001963 | FALSE | FALSE | 930 | Ace |
| ENSMUST00000061516 | FALSE | FALSE | 160 | Fpr1 |
| ENSMUST00000036387 | FALSE | FALSE | 1091 | S100b |
| ENSMUST00000029078 | FALSE | FALSE | 680 | Car2 |
| ENSMUST00000102715 | FALSE | FALSE | 1474 | Stk39 |
| ENSMUST00000114261 | FALSE | TRUE | 180 | Paxx |
| ENSMUST00000040647 | FALSE | FALSE | 3254 | Fgf1 |
| ENSMUST00000034777 | FALSE | FALSE | 347 | Calml4 |
| ENSMUST00000037287 | FALSE | FALSE | 685 | Cdkn1c |
| ENSMUST00000035196 | FALSE | FALSE | 952 | Hemk1 |
| ENSMUST00000068545 | FALSE | FALSE | 383 | Sugct |
| ENSMUST00000109375 | FALSE | FALSE | 883 | Elmo3 |
| ENSMUST00000127247 | 0 | TRUE | 3497 | Tmem229a |
| ENSMUST00000103108 | FALSE | FALSE | 374 | Wnk4 |
| ENSMUST00000030922 | FALSE | FALSE | 723 | Prkcz |
| ENSMUST00000074839 | FALSE | FALSE | 177 | Ear2 |
| - | - | - | - | |
| ENSMUST00000098331 | FALSE | FALSE | 1315 | Cpeb1 |
| ENSMUST00000039318 | FALSE | FALSE | 4163 | Cdc14b |
| ENSMUST00000159175 | FALSE | FALSE | 178 | Ear10 |

| | | | | |
|---|---|---|---|---|
| ENSMUST00000024594 | FALSE | FALSE | 566 | Agpat4 |
| ENSMUST00000020003 | FALSE | FALSE | 489 | Fam184a |
| ENSMUST00000066489 | FALSE | FALSE | 666 | P3h4 |
| ENSMUST00000075639 | FALSE | FALSE | 51 | Lrmda |
| ENSMUST00000106372 | FALSE | FALSE | 539 | Sult1a1 |
| ENSMUST00000118522 | FALSE | FALSE | 1024 | Paxbp1 |
| - | - | - | - | |
| ENSMUST00000032174 | FALSE | FALSE | 950 | Klf15 |
| ENSMUST00000095446 | FALSE | FALSE | 314 | Adamtsl5 |
| ENSMUST00000049544 | FALSE | TRUE | 4464 | Ccdc141 |
| ENSMUST00000055223 | 0 | FALSE | 0 | Dnajb2 |
| ENSMUST00000019721 | FALSE | FALSE | 2098 | Pdk4 |
| ENSMUST00000164886 | FALSE | FALSE | 1024 | Pilrb2 |
| - | - | - | - | |
| ENSMUST00000040320 | FALSE | FALSE | 3779 | Micall1 |
| ENSMUST00000018765 | FALSE | FALSE | 947 | Nmnat1 |
| ENSMUST00000003620 | FALSE | TRUE | 451 | Prodh |
| ENSMUST00000165999 | 0 | TRUE | 0 | Cyp4f17 |
| - | - | - | - | |
| ENSMUST00000079413 | FALSE | FALSE | 1953 | Fkbp5 |
| ENSMUST00000034244 | FALSE | FALSE | 1196 | Tmem38a |
| ENSMUST00000036418 | FALSE | FALSE | 129 | Ciart |
| ENSMUST00000006828 | FALSE | FALSE | 311 | Aplp1 |
| ENSMUST00000049389 | | | | |
| ENSMUST00000025965 | | | | |
| ENSMUST00000029440 | | | | |
| ENSMUST00000065976 | | | | |
| ENSMUST00000134825 | | | | |
| ENSMUST00000028826 | | | | |
| ENSMUST00000113306 | | | | |
| ENSMUST00000115390 | | | | |
| ENSMUST00000021610 | | | | |
| ENSMUST00000020400 | | | | |
| ENSMUST00000121390 | | | | |
| ENSMUST00000019101 | | | | |
| ENSMUST00000021784 | | | | |
| ENSMUST00000070518 | | | | |
| ENSMUST00000048820 | | | | |
| ENSMUST00000041105 | | | | |
| ENSMUST00000089236 | | | | |
| ENSMUST00000020779 | | | | |
| ENSMUST00000197515 | | | | |
| ENSMUST00000169803 | | | | |
